# Experimentally unsupervised deconvolution for light-sheet microscopy with propagation-invariant beams

**DOI:** 10.1101/2021.05.26.445797

**Authors:** Philip Wijesinghe, Stella Corsetti, Darren J.X. Chow, Shuzo Sakata, Kylie R. Dunning, Kishan Dholakia

## Abstract

Deconvolution is a challenging inverse problem, particularly in techniques that employ complex engineered point-spread functions, such as microscopy with propagation-invariant beams. Here, we present a deep learning method for deconvolution that, in lieu of end-to-end training with ground truths, is trained using known physics of the imaging system. Specifically, we train a generative adversarial network with images generated with the known point-spread function of the system, and combine this with unpaired experimental data that preserves perceptual content. Our method rapidly and robustly deconvolves and super-resolves microscopy images, demonstrating a two-fold improvement in image contrast to conventional deconvolution methods. In contrast to common end-to-end networks that often require 1,000–10,000s paired images, our method is experimentally unsupervised and can be trained solely on a few hundred regions of interest. We demonstrate its performance on light-sheet microscopy with propagation-invariant Airy beams, including in calibration beads, oocytes, preimplantation embryos, and excised brain tissue, as well as illustrate its utility for Bessel-beam LSM. This method aims to democratise learned methods for deconvolution, as it does not require data acquisition outwith the conventional imaging protocol.

## Introduction

Deconvolution is a well-known problem in many fields of engineering and science (1–3). Specifically in computational imaging, the problem of deconvolution is one of the accurate reconstruction of the imaged object from data encoded by the imaging system. This encoding, for instance through the use of structured light fields and engineered point-spread functions (PSF), can convey much greater information about the imaged object than is possible with conventional imaging (4). Propagation-invariant beams, in particular, have extended the field of view and resolution (5, 6), and have demonstrated deeper penetration into scattering tissues (7, 8). These methods have pushed the envelope of many important trade-offs, such as those between the resolution, contrast, field of view, imaging speed and photo-damage.

Specifically, Airy and Bessel beams have enabled rapid multiscale imaging at exceptional resolution in light-sheet microscopy (LSM) (5, 9). These beam shapes can be generated using accessible optical components and implemented in simple optical geometries (5, 10, 11). Despite their superior performance (5, 9), they are yet to be broadly adopted by the microscopy community due to the practical challenges of deconvolution. This is because these PSFs are often burdened with side-lobe structures, which challenge contemporary deconvolution algorithms, more-so than a Gaussian blur, especially in the presence of poor signal-to-noise (SNR), scattering and speckle (3). As a consequence, LSM with propagation-invariant beams often fails to produce image contrast that exceeds conventional Gaussian-beam LSM at focus.

To overcome these major barriers, in this paper, we introduce experimentally unsupervised deconvolution based on deep learning that is capable of robustly and rapidly deconvolving and super-resolving microscopy images with arbitrary PSFs. Recently, deep learning has emerged as a method to solve a variety of inverse problems in microscopy (4, 12). For instance, deep learning has enabled super-resolution from single images (13, 14).These networks are typically trained using a supervised end-to-end approach, where the network learns directly the transformation from low-resolution inputs to high-resolution ground truths, which are paired and coregistered experimentally using higher-resolution acquisition modes or even across modalities (14), typically requiring several 1,000s to 10,000s of images. Image enhancement methods have further been extended to 3D using efficient dense residual network architectures (15, 16). Whilst such networks would likely be capable of addressing deconvolution of PSFs from propagation-invariant beams, the onus is placed on the user to acquire sufficient data matching structured PSFs with high-resolution ground truths via modifications to imaging instruments and extended experiments. Further, many such methods learn, non-discriminately, the inversion of the physical system and the properties of the samples contained in the training set (4). This may lead to poor generalisation and, thus, poor performance when a trained network is applied to samples outwith the training parameters (17). Thus, the user is required train bespoke networks for each system and sample imaged, which is a present and major barrier to broad adoption.

Recent networks have also addressed the onus of extensive ground truth matching. For instance, unpaired high-resolution ground truths were digitally degraded to estimate low-resolution network inputs, such that subsequent under-sampled acquisitions may be reconstructed closer to the original image quality (16), enabling thigh-throughput imaging. Alternatively, super-resolution ground truths could be generated computationally using conventional algorithms, such as SRRF (16). In similar ‘model-based’ approaches, the output of deconvolution using a back projection algorithm was used to train a network to speed up processing (18), and high-resolution *xy* cross-sections in microscopy have been used as a ground truth target to deconvolve *xz* sections to achieve isotropic volumes (19). In these methods, deep learning has provided an exceptional speed-up and has reduced data requirements. However, these approaches still either require experimental ground truths or are unlikely to exceed the quality of the original numerical algorithms used to train them.

Experimentally unsupervised networks, trained using simulated ground truth data, have emerged for image recovery problems (12). Such methods have been instrumental in phase unwrapping (20), where ground truths are intractable to obtain experimentally, and have recently enabled the reconstruction of structured illumination microscopy (21). CARE microscopy has proposed the use of simulated ground truths that resembled expected sample content, such as microtubules and secondary granules, showing superior deblurring to conventional deconvolution methods (13). In this paper, we extend on this work, and illustrate that a generalised network for deconvolution of engineered PSFs can be trained with good performance without any measurement or prediction of a ground truth. We achieve this by training a network that is informed by physics priors, *i.e*., a priori knowledge of the physics of the imaging system, and minimising content priors, such as the predictions of what the image content should look like (4, 22). This physics-informed learning has emerged to reduce the need for experimental training data and to direct training towards generalisation that is agnostic of the samples being imaged (4, 12), bridging the gap between conventional algorithms and data-driven end-to-end networks. In this paper, we follow the inspiration of such networks and make use of three key priors: (1) we make use of the known PSF of the imaging system to generate simulated paired data that is consistent with images that can be generated by the experiment; (2) the generated images are controlled in their spatial and spatial-frequency content, thus, provide control over the sparsity expected in the network outputs; and (3) we use experimental images to guide reconstruction towards the perceptual quality and power spectral content expected in the imaging system.

Importantly, our method is entirely experimentally unsupervised, i.e., it does not require experimental ground truths, and may be trained rapidly on a single light-sheet volume, including the one that is desired to be deconvolved. Our network is a simple approach to deconvolution with a known PSF that mirrors the utility of common algorithms such as Richardson-Lucy deconvolution. We demonstrate that the learned approach is superior to iterative deconvolution methods in its ability to achieve a more symmetric deconvolved PSF, a 3–5 dB% increased peak signal-to-noise ratio, and can be performed rapidly at 0.2 s per widefield image. Further, we illustrate that our method is robust to noise and is 3-fold more resistant to model mismatch, which is advantageous in the presence of misalignment or aberrations. We validate our method experimentally on LSM using an Airy beam in calibration beads, autofluorescent mouse oocytes and embryos, and excised mouse brain tissue. We supplement this validation with demonstrations in multiphoton LSM with the commonly used Bessel beam for developmental biology. The method generalises well, learning primarily the inversion of the imaging operator, which we demonstrate by training only a single network for each beam shape, and applying it to all of the samples we subsequently image. Our approach to deep learning makes the use of structured light fields and deconvolution highly accessible as it does not require acquisition outwith the conventional imaging protocol. Towards this, we illustrate its use in an openly available dataset, and provide the deep learning code as open source to facilitate broad evaluation of this approach among the imaging community.

## Results

### Network design

Figure 1 illustrates the architecture of our method under training in the context of the LSM setup. The network architecture is described in detail in the Methods Section, and briefly summarised here. We perform deep learning based on a generative adversarial network (GAN) (23) and several tuned training loss functions (TLFs). The inverse problem, i.e., the transformation between an image encoded with a structured PSF (LR) to a high-resolution estimate of the ground truth, is learned using a Generator (G) network, based on a 16-layer residual network (ResNet) (24) (Supplementary Notes 1 and 4). Concurrently, another Discriminator (D) network (PatchGAN (25)) is trained to identify network outputs (DL) and real ground truths (GT). We have found that this adversarial training leads to the learning of strong image features that are consistent with the GT and that are difficult to capture using pixel-wise TLFs (Supplementary Note 3), which is consistent with previous investigations (26).

**Fig. 1.**
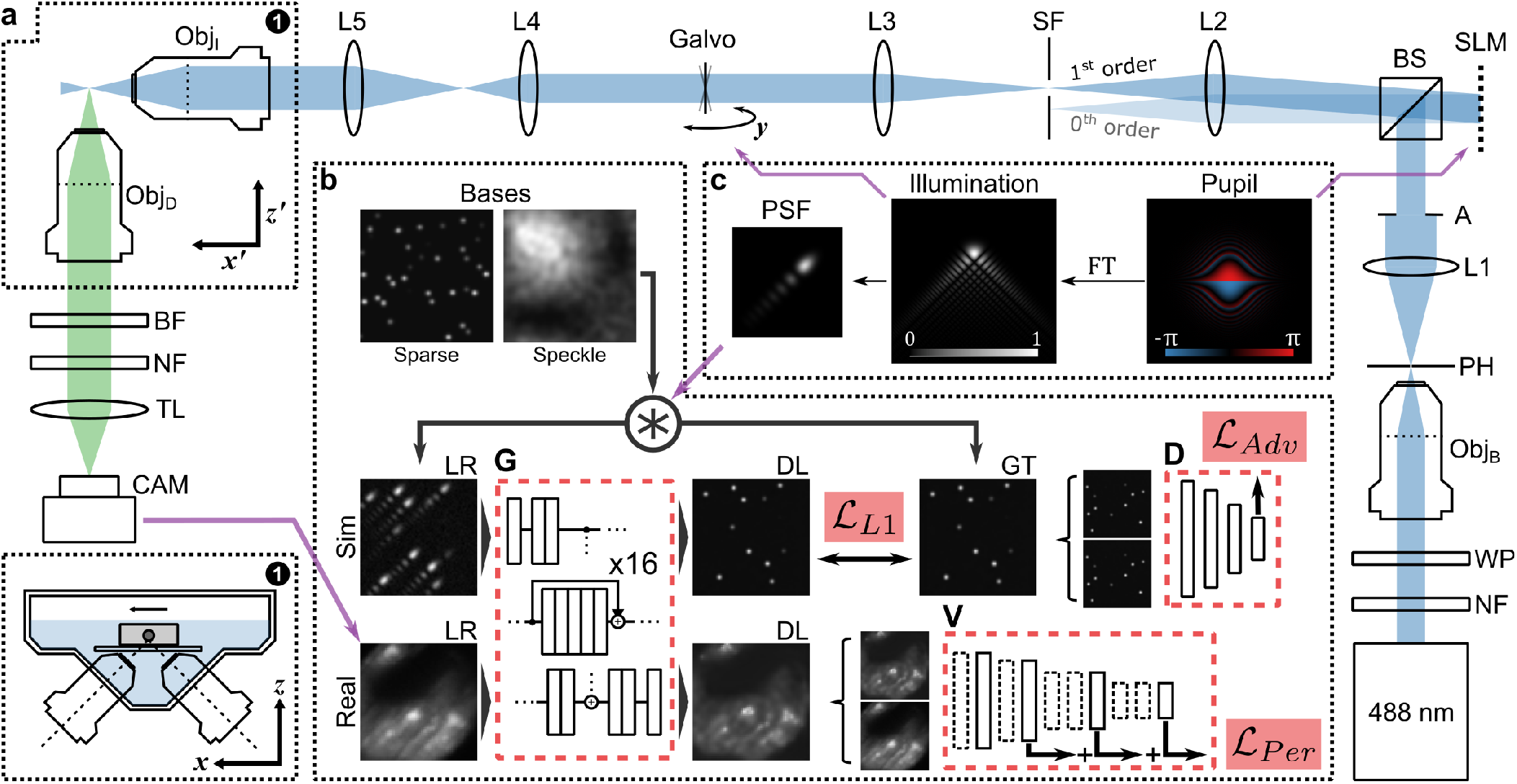
Light sheet microscopy system and physics-informed learned deconvolution training. (a) Optical geometry of the system. Insets marked by (1) show the optical and physical positioning of the sample, which are denoted using the *x′ z′* and *xz* coordinates, respectively. (b) Deep learning method, consistent with (c) the physics of beam shaping using a spatial light modulator (SLM). Obj B, I, D: objectives for beam expansion, illumination and detection respectively. A 488 nm laser source is filtered using a notch filter (NF), half wave plate (WP), pinhole (PH) and aperture (A). L1-5: lenses. BS: beamsplitter. The SLM shapes the illumination field in diffraction mode to match the pupil function in (c), which is filtered using a spatial filter (SF) and scanned as a light sheet using a galvonometer driven mirror. Sample fluorescence filtered using a bandpass filter (BF) and a notch filter (NF), and imaged using a tube lens (TL) onto the camera (CAM). Deep learning learns from simulated images generated from sparse and speckle bases and the known PSF, and the real images captured by the CAM. During training, the generator (G) learns from a combination of *L*_1_ pixel loss, adversarial loss from the discriminator (D) and the perceptual loss from the perceptual loss network (V).

Figure 1(a) illustrates the LSM geometry (see Methods for details). An Airy light field is generated using a nematic spatial light modulator (SLM) in diffractive mode, encoding the appropriate cubic phase profile (the Fourier transform of the Airy function). The SLM enables precise control of the field intensity and phase in illumination. The desired pupil phase function (Fig. 1(c)), i.e., the spatial Fourier transform of the beam cross-section at focus, is projected onto the SLM (Supplementary Note 14). In this way, it is trivial to estimate the PSF of the LSM by combining the Fourier transform of the pupil, evaluated at the spatial coordinates established by the various lenses, and the diffraction-limited Gaussian spot size of the detection. Here, we train and perform deconvolution in 2D, in the cross-sectional plane of the illumination and detection objective (Fig. 1(a), inset (1)). This plane includes the PSF structure of the propagation-invariant light fields, and corresponds to the lowest resolution plane of the system. In this geometry, training and deconvolution in 2D is a close approximant to a 3D model, whilst remaining computationally accessible to consumer hardware (Supplementary Note 8).

During a single training iteration, a batch of simulated images consistent with the LSM is fed into G. These images comprise two domains: a sparse collection of points, and speckle (Fig. 1(b)). Network inputs and their ground truths are formed, respectively, by convolving the images with the system PSF and a Gaussian spot set to 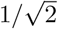 of the diffraction limit of the objectives. The sparse points provide strong gradients for the network to learn the PSF shape. Speckle is a Fourier-transform pair to a uniformly distributed random phase and a Gaussian spectral amplitude. Thus, speckle images encode a wide spatial-frequency content with low sparsity. This pushes the network to prioritise physical learning (i.e., the transformation by the PSF) over content learning (i.e., the expectation of what microscopy images should look like), which benefits generalisation. During training, the network outputs are compared to the ground truths using the Discriminator (adversarial loss), and using L1-norm error (pixel loss). Concurrently, a random selection of experimental images are processed by G. The inputs and outputs of the network are compared using a pre-trained perceptual loss network (VGG-16 (27)), V (Fig. 1(b)), which focuses on the conservation of salient image content (28) (Supplementary Note 2). The addition of this TLF leads to the preservation of the power spectral density of experimental images, which is an important prior (29), and is effective at eliminating network artefacts. The combined loss of G is the linear combination of the adversarial, pixel and perceptual losses. The Discriminator is trained using conventional means (25), and training is alternated evenly between G and D.

Our method and architecture results in a robust and generalised model. Importantly, and in contrast with most learned image enhancement methods (12), we demonstrate this by training only a single network for each beam shape using only one experimental low-resolution volume of mouse embryos (~200 images) and no experimental ground truths. We apply this network, non-discriminately, to a wide array of samples, namely fluorescent beads samples, other embryos, oocytes, turbid brain tissue and zebrafish embryos.

### Learned deconvolution

Here, we illustrate that DL can achieve superior performance to numerical methods, specifically the well-known Richardson-Lucy (RL) algorithm (30, 31) and RL with added total variation regularisation. This is because, firstly, we forego the common assumption that noise is a uniform and randomly distributed Poisson process (3). Secondly, neural networks naturally use sparsity in images to improve reconstruction (32). Finally, as we will demonstrate, our network is less prescriptive than RL deconvolution and can tolerate greater model mismatch, for instance, when the theoretical and experimental PSFs differ. This means that our network is likely to preserve its good performance even if the microscope is not perfectly aligned.

First, we quantify performance with simulated data on several propagation-invariant beam types. Figure 2(a) evaluates deconvolution in images simulated by convolving a blastocyst embryo GT with PSFs generated by the Airy, Bessel and Gaussian beams. The networks were trained solely on a collection of sparse and speckle bases with no knowledge of the blastocyst images. The generation of the various beam shapes is described in the Methods section and in Supplementary Note 14. Briefly, the Airy beams are generated by a cubic phase in the pupil plane, with a scale-invariant *α* parameter that corresponds to the gradient of the cubic function. The Bessel beams are formed by a ring in the pupil plane, parameterised by a ring’s radius and cross-sectional width. We later focus on the Airy and Gaussian beams in the experiment as they are readily generated by our LSM configuration.

**Fig. 2.**
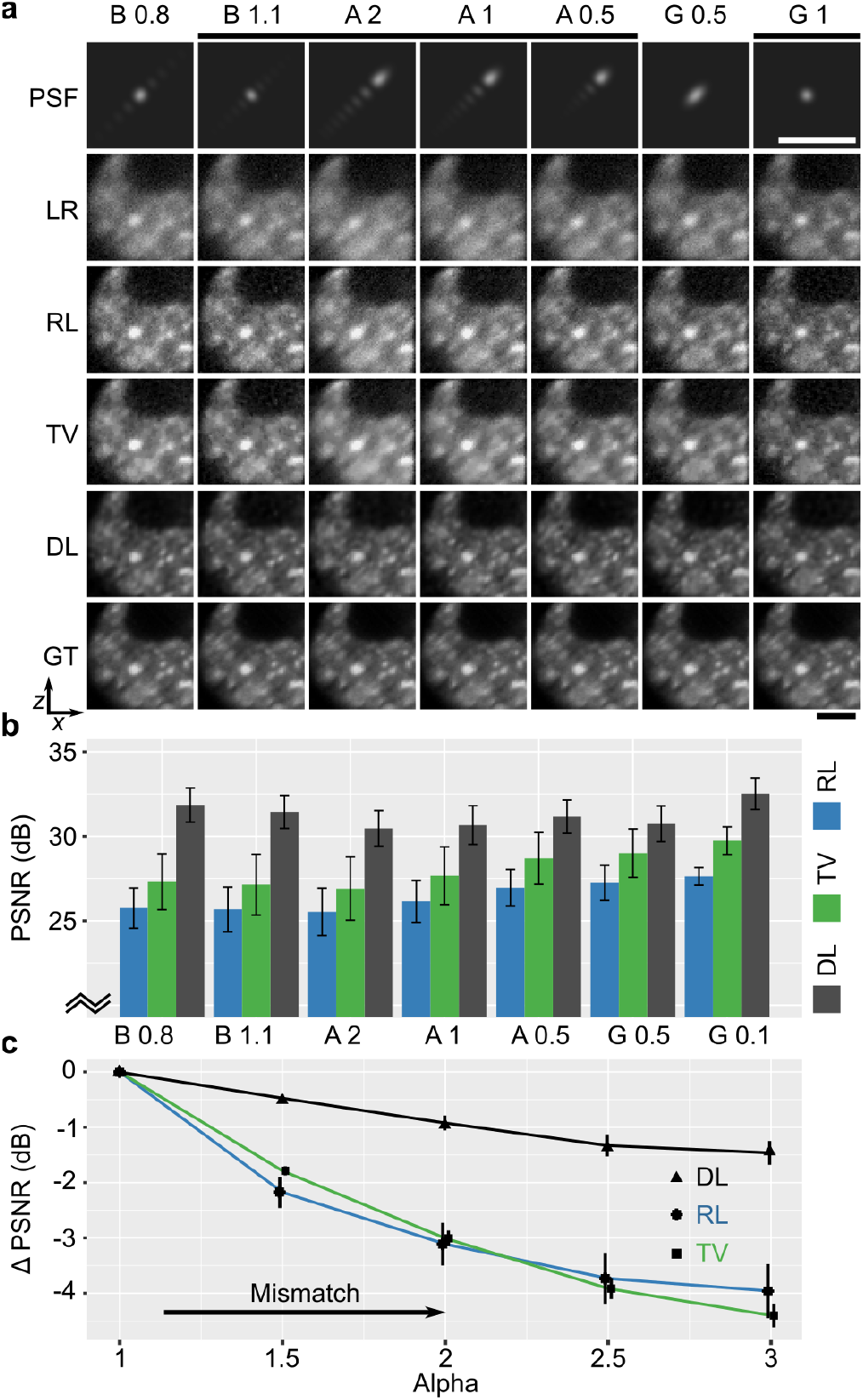
Comparison of deep learning (DL), Richardson-Lucy (RL) deconvolution and RL deconvolution with total variation (TV) regularisation in simulated data. (a) Deconvolution of low-resolution noisy images (LR) simulated using beams (B: Bessel; A: Airy; G: Gaussian) visualised by their point-spread function (PSF) (parameters listed in Methods) using RL, TV and DL algorithms and compared to the ground truth (GT). Scale bar is 20 *μm.* (b) Mean peak signal-to-noise ratio (PSNR) of RL, TV and DL deconvolution for each beam shape evaluated in 240 simulated images (Error bar is standard deviation). (c) Relative change in PSNR in deconvolution of an Airy beam due to model mismatch. RL, TV and DL assume that the Airy *α* =1; however, LR images were simulated with *α* in [1,3].

From Figure 2(a) it is evident that when using propagation-invariant beams, light-sheet images (LR) exhibit substantial side lobes. Here, the LR images include randomly distributed Gaussian (thermal) and Poisson (shot) noise. The conventional RL algorithm is able to reduce side lobes and present a spatially consistent image to the GT. However, RL-deconvolved images fail to present a visual quality matching that of the GT or the high-resolution Gaussian beam. Despite losing the structure, side lobes persist as elevated background intensities. We further include a common implementation of RL-deconvolution with total variation regularisation (TV), which exhibits a similar performance to RL but with reduced high-frequency noise. The improvement from DL is also consistent when compared to other direct and iterative deconvolution algorithms (3), which we illustrate in Supplementary Note 6.

Beam shapes marked by a black bar in Figure 2(a) carry similar spatial resolution. DL is capable of deconvolving LR images much closer to the GT, eliminating the side lobes and presenting a Gaussian-like PSF. This is especially clear in simulated images of beads in Supplementary Note 5. Figure 2(b) quantifies the performance of these methods using the peak signal-to-noise ratio (PSNR) over 240 simulated images. It is evident that DL substantially outperforms RL and TV deconvolution, achieving a 30–33 dB PSNR, whilst RL and TV range in 25–30 dB (similar to other deconvolution algorithms in Supplementary Note 6.). We note that a 3 dB change in PSNR is approximately a two-fold change in noise. DL further outperforms RL consistently with increasing noise and with added background fluorescence (Supplementary Note 7). It is important to note that convergence of RL iterations leads to high image noise and early truncation is often used as a spatial regulariser. Here, and for all further RL deconvolution, we have truncated RL to 6 iterations specifically to maximise the PSNR parameter for a best-case comparison. Further, we used a TV regularisation parameter of 0.001 for the most visually compelling recovery.

A major challenge in deconvolution and in deep learning with priors is model mismatch (4). Here, this arises when the theoretical PSF differs from that of the imaging system, for instance, due to misalignment or aberrations. Figure 2(c) illustrates the robustness of our method to model mismatch. Specifically, we have generated LR images corresponding to Airy beams with *α* values in the range of 1 to 3. However, to deconvolve the images, we have used a network trained on an Airy beam with an *α* = 1. Similarly, we have used a single Airy (*α* = 1) PSF as a reference for RL and TV deconvolution. The matched training and testing case of *α* = 1 was used as a reference. Figure 2(c) shows the relative degradation in the PSNR (to the known GT) as greater mismatch is introduced. It is clear that DL tolerates much greater mismatch than RL and TV deconvolution, on the order of 3-fold improvement (2.7–3.8 times) to the error metrics. While blind deconvolution methods may overcome some issues of mismatch, we have observed that blind deconvolution struggles to converge when the PSF is structured (*i.e.,* it is not monotonically decreasing from a central peak) and when images are not spatially sparse, which is consistent with previous observations (3).

Figure 3 extends the demonstration of our DL method to experimental data of 200-nm beads. Using the SLM, we readily generate a Gaussian illumination (Fig. 3(a)), and compare it to the Airy beam with *α* parameters of 0.5, 1 and 2, respectively (Figs. 3(b–d)). The red bar indicates the Rayleigh range of each beam shape, which corresponds to the field of view available for LSM. The acquired LSM images are labelled as LR, and are deconvolved using DL and RL deconvolution. Here, we exclude TV regularisation for clarity due to its emphasis on sharp discontinuities. For each beam shape, the network was trained on one acquisition of tissue data (see Methods) and applied to all experimental data included in this work. The 200-nm beads are sub-diffraction limit, thus, represent the system response or the PSF. We can see that DL offers superior image quality to RL deconvolution. This is clear in the transverse profile of the beads marked by the red box. The Airy function can be seen in the LR profiles. The RL deconvolution removes the structure of the side lobes, but fails to substantially reduce the envelope, leading to an asymmetric PSF. The superior performance of DL is evident by the sharp and more Gaussian-like PSF.

**Fig. 3.**
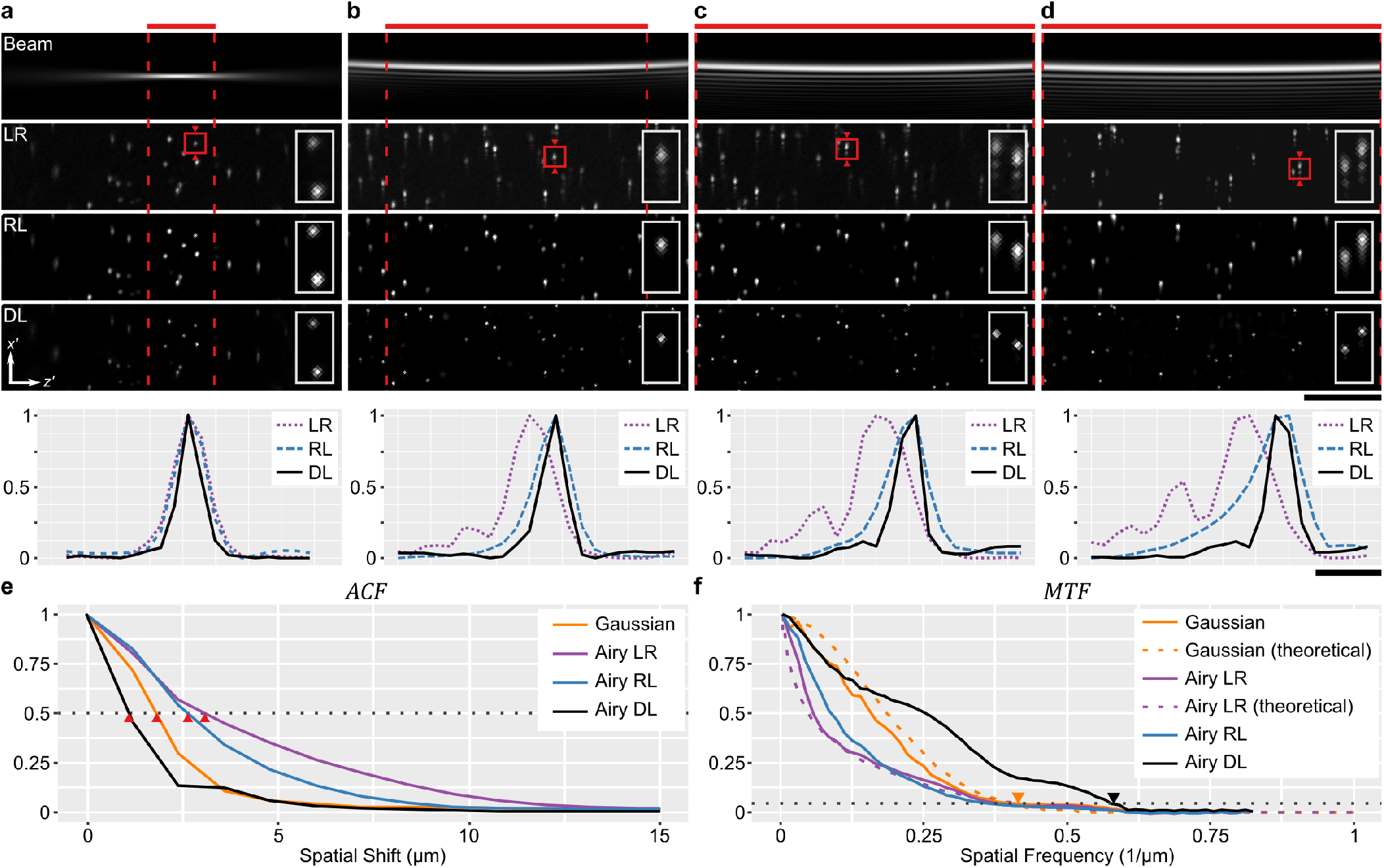
Learned deconvolution in beads using a (a) Gaussian, (b) Airy (*α* = 0.5), (c) Airy (*α* = 1), and (d) Airy (*α* = 2) beam. (a–d) Show the illumination beam and the corresponding acquired LSM image (LR), which is deconvolved using Richardson-Lucy (RL) and deep learning (DL) methods. Insets and profiles correspond to the bead marked by a red square. Red dashed lines mark the Rayleigh range of each beam. Scale bars are 50 *μm* for images and 5 *μm* for plots. (e) Autocorrelation function (ACF) and (f) Modulation transfer function (MTF) with respect to depth of the Gaussian and Airy (*α* = 1) LSM images.

In addition to decoding the structure, DL leads to a superresolution of individual beads beyond the diffraction limit of the light field. We can quantify this performance over many beads in the field of view using the autocorrelation function (Fig. 3(e)), which represents the shift invariance and relates to the depth or axial LSM resolution. From the Wiener–Khinchin theorem, we know that the ACF of intensity is a Fourier pair to the power spectral density, *i.e.,* the square of the modulation transfer function (MTF). Figure 3(f) shows the MTF of the Gaussian LSM and the Airy (*α* = 1) beams compared to their theoretical MTF, as well as the RL and DL deconvolutions. We can see that the MTF in the acquired images match theory well (dashed lines). The dotted line marks the 5% threshold of the MTF, which corresponds approximately to the full-width at half-maximum (FWHM) of the PSF. We note that the intersection of the Gaussian and Airy beam MTFs at the 5% threshold indicates their capacity to carry similar high-frequency content. However, this information is not well conveyed by the Airy images in the spatial coordinate space as illustrated by the widened ACF. As expected, RL deconvolution decodes Airy images (LR) to improve the ACF width with little benefit seen in the MTF. Remarkably, DL expands the MTF beyond that of the Gaussian beam, and extends the crossing of the 5% threshold by a factor of 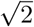. The improvement in the resolution is consistent with the simulation of the HR training data at 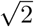 of the LR images. In principle, it is possible to tailor the resolution improvement based on the choice of the priors. However, larger resolution improvements are met with lowered contrast at lower spatial frequencies. Our choice of a 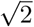 improvement was motivated empirically by visually inspecting the network outputs. This improved performance is also evident by the rapidly decreasing ACF. The FWHM resolution was quantified as 2.6 *μm* in the Gaussian and Airy images, and 1.7 *μ*m with the use of DL. This exemplifies the power of deep learning as a method to solve inverse problems that leverages sparsity, learned noise statistics and the imaging operator to both decode information and extend the bandwidth limit.

DL has provided the Airy LSM a 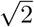 enhancement in resolution over a substantially wider FOV of up to 372 *μ*m vs. the 27-*μ*m FOV of the Gaussian beam, a greater than 13 fold improvement. The accessible and rapid inference of DL, the wide FOV and the ease of generating propagation-invariant beams confers exemplary potential to multiscale imaging applications. To demonstrate this, we explore our DL method in mouse oocytes and embryos, and in excised mouse brain tissue.

### Oocytes and embryos

Rapid minimally invasive imaging of embryos is important for diagnostics and may aid in the monitoring of embryo development following in vitro fertilisation (IVF) (33, 34). LSM is particularly promising in this area due to its rapid volumetric imaging, reduced photodamage and low cost compared to confocal microscopy (33). Our DL-assisted Airy LSM is capable of performing cross-sectional imaging of the entire depth of an embryo (~180 *μ*m) in one shot, i.e., within a single acquisition on the camera. Further, it is able to convey high-resolution morphological and functional information from autofluorescence.

Figure 4 demonstrates the performance of DL deconvolution in mouse blastocyst-stage embryos. The blastocyst is the final stage of preimplantation embryo development, and comprises two sub-populations of cells: the inner cell mass (ICM) which forms the fetus, and the trophectoderm (TE) which leads to the development of all extra embryonic tissues, including the placenta (35). Figure 4(a) shows a crosssection through a blastocyst imaged with an Airy (*α* = 1) beam and deconvolved using RL and DL methods. The same region was imaged using a Gaussian beam (G). We can see that DL readily decodes the Airy PSF from the LR image, and matches and exceeds the image quality presented by the Gaussian LSM. We illustrate this further by taking a profile marked by the red triangles. In Figure 4(b), we can see that quantitatively the DL profile matches that of the Gaussian GT, while the RL fails to match the relative intensity profile. It is important to mention that the LR profile does not match that of G. This is because the Airy PSF structure (here, extending top-right to bottom-left in the images) has distinct oscillating side lobes. Thus, the local intensity varies non-monotonically with distance from the scattering object. Deconvolution must be performed on such images to decode the underlying quantitative fluorescence distribution, which in our case is done well with DL compared to RL. This can be further evidenced in similar cross-section in Supplementary Note 9. Using Fourier analysis (Supplementary Note 10), the spatial resolution was quantified as 5.0 *μ*m in LR, 4.9 *μ*m in RL, 3.8 *μ*m in G, and 3.4 *μ*m in DL images.

**Fig. 4.**
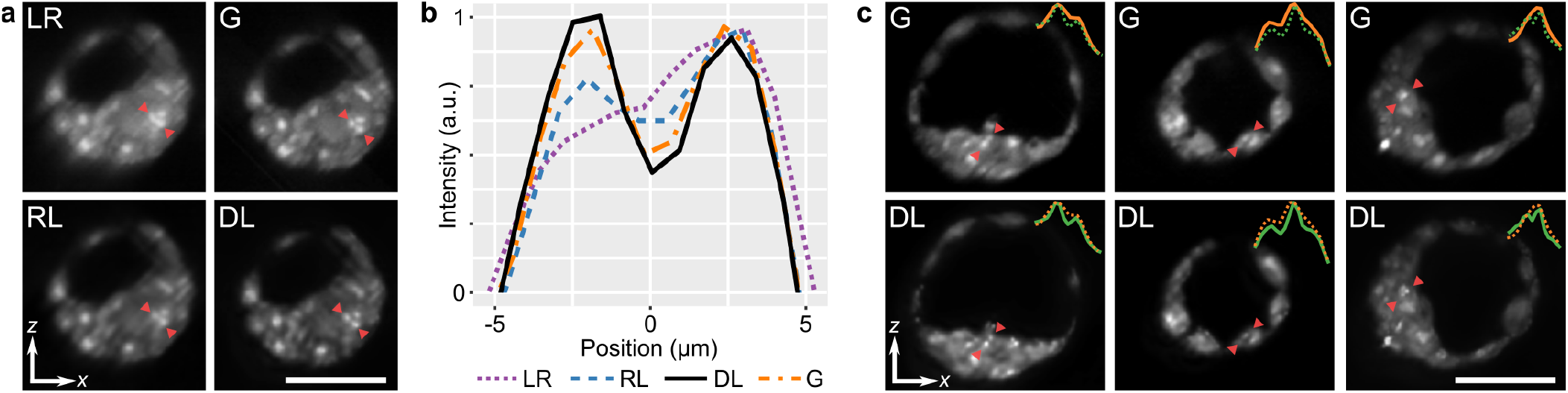
Cross-sectional images of mouse blastocyst-stage embryos. (a) Airy LSM image (LR) is deconvolved with Richardson-Lucy (RL) and deep learning (DL) deconvolution, and compared with a Gaussian LSM (G). (b) Intensity profiles marked by the red arrows in (a). (c) Cross-sections of blastocyst-stage embryos imaged with a Gaussian LSM (G) and compared to an Airy LSM with deep learning deconvolution (DL). Profiles marked by the red triangles are displayed as insets. Scale bar is 50 *μ*m.

We now focus on demonstrating the advantage of using the Airy beam LSM with deep learning deconvolution compared to the conventional Gaussian LSM, which represents the most popular embodiment of LSM. Figure 4(c) shows several cross-sections from different blastocyst-stage embryos imaged with a Gaussian LSM (G) compared to matching regions imaged with an Airy LSM with DL, hereafter referred to as ‘Airy DL’ for succinctness. Visually, we see an improvement in the contrast and resolution in the DL images. Line profiles marked by the red triangles and inset into the figures further illustrate that DL is able to distinguish autofluorescent features with a higher spatial resolution. We attribute these bright features to active mitochondria performing high levels of metabolic activity and, thus, generating higher amount of intracellular fluorophores, such as flavin adenine dinucleotide (FAD) (36). This is consistent with the excitation and emission filters in our LSM setup. Resolution, intensity and count of these features may be an important metric for IVF success (34). It is clear that the added resolution may assist the counting of individual sharp features, and the FOV in depth enabled doing so in a single camera snapshot.

Figure 5 compares a Gaussian LSM to Airy LSM with DL in mouse cumulus oocyte complexes (COCs). COCs comprise of an oocyte at the centre, surrounded by much smaller cumulus cells. Imaging of the COC may provide an opportunity to assess the health of the oocyte prior to IVF (36). The larger field of view encompassing multiple COCs emphasises the major advantage of using an Airy beam, namely, the extended depth of field (DOF). Figure 5(a) and (b) shows the widefield cross-sections and Fig. 5(c) emphasises the image enhancement of Airy DL over conventional Gaussian LSM in the identification and inspection of cumulus cells in COCs. These sections are maximum intensity projections over a 50 *μm* range to emphasise COC morphology. Figure 5(a) and (b) mark the focus and DOF of the illumination beams by the red triangle and bar, respectively. It is clear that much of the Gaussian cross-sections are out of focus. Airy DL is able to maintain the COC volume in the focal region and demonstrates clear morphological features and enhanced resolution, both in and out of focus. In the Figure 5(c), we can see the clear demarcation of cumulus cells, even in regions well beyond the focus available to the Gaussian beam. In focus, we have quantified the resolution (Supplementary Note 10) to be 4.3 *μ*m for LR, 3.9 *μ*m for Gaussian, and 3.1 *μ*m for Airy DL images.

**Fig. 5.**
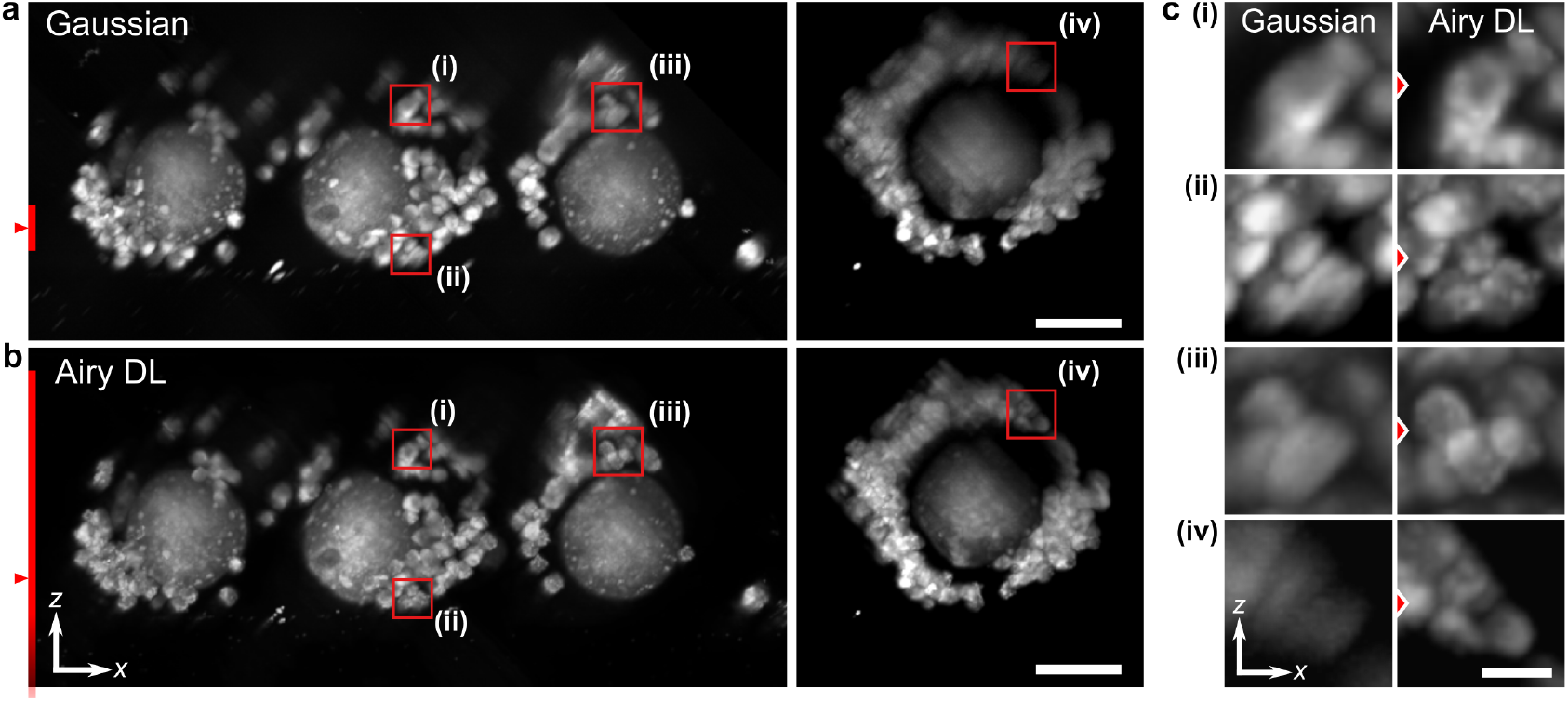
Cross-sectional LSM images of mouse cumulus oocyte complexes (COCs) with a Gaussian and Airy beams with deep learning deconvolution. (a,b) Widefield crosssectional maximum intensity projections (range: 50 *μm*). (c) Zoomed-in regions of cumulus cells (i–iv) emphasising the enhancement of Airy DL over Gaussian LSM from corresponding regions in (a,b) marked by red rectangles. The red line on the left of (a,b) shows the theoretical DOF of the Gaussian and Airy beams. The red triangle marks the focal position. Scale bars are (a,b) 50 *μ*m and (c) 10 *μ*m.

The volumetric and widefield imaging capacity of Airy DL LSM is further detailed in Supplementary Note 11. It shows enface (*xy*) sections corresponding to Fig. 5 and illustrates an oocyte arrested at the metaphase II stage of meiosis. During metaphase, the chromosomes are aligned and held in place by the meiotic spindle along a plane termed the metaphase plate. This can be seen by the white line that separates two darker spherical regions that depict the spindle barrel. This morphology is particularly prominent in the Airy DL images, and can be confirmed for all oocytes by inspecting the volume (see Data Availability). This is an essential event for an oocyte undergoing nuclear maturation, necessary for fertilisation. As described for the blastocyst-stage embryo, the small bright regions within the oocyte are indicative of metabolic activity of mitochondria through FAD autofluorescence. The multitude of morphological features enhanced by Airy DL over a wide FOV and depth range underlines its potential for label-free and low-phototoxic imaging for embryo health and IVF success. Revealing the source and implications of these markers would be of considerable interest for future bioimaging studies.

### Brain

LSM is a powerful emerging technique for neuroscience (37). It is well positioned to address multiscale imaging at depth, combining imaging at the resolutions and timescales of synapses, over tailored fields-of-view corresponding to whole brains of model organisms or regions of pathogenesis. However, penetration depth is a present challenge, especially in single-photon modalities. The Airy light field has several properties that are of great benefit in this area. Beyond its propagation invariance, Airy light fields are self-healing. If a portion of the field is blocked or scattered, the beam cross-section will reform into its expected Airy function. This has led to several observations that Airy light fields penetrates on the order of 30% deeper into tissues (7, 38).

Towards this, we demonstrate our technique in excised mouse brain that expresses enhanced YFP in parvalbumin-positive (PV+) neurons. Figure 6(a) and (b) show enface (*xy*) sections with depth taken from cortical areas, including the hippocampus. These regions were selected from a larger acquired volume, indicated in the cross-sectional *xz* image (Fig. 6(c)). To the right of each *xy* image are zoomed-in regions corresponding to the areas marked by the white dashed rectangles. The *xy* sections visualise the imaging performance in depth, and focus on regions of interest selected by the plane of brain dissection. Because PV+ neurons provide somatic inhibition in the hippocampus (39) and the neocortex (40), we hypothesise that axonal boutons in perisomatic regions (around the cell body) clearly mark PV+ neurons in Airy DL images as bright rings (red asterisks). While PV+ neurons are clearly seen, axonal terminals (bright small spots) around the cell bodies (large dark circles) are also visible in other neurons in lower number. Close to the surface, the enhancement in contrast and resolution of the Airy DL, for instance when inspecting the areas marked by the red asterisk, matches that seen in the blastocysts (Fig. 4). We have quantified the resolution (Supplementary Note 10) to be 7.1 *μ*m for LR, 5.3 *μ*m for Gaussian, and 4.5 *μ*m for Airy DL images. Image quality in the Gaussian LSM rapidly deteriorates with depth, showing substantial blurring and loss of contrast. However, image quality is preserved in depth with Airy DL LSM, distinguishing sharper fluorescent features. This is particularly evident at 52 *μm* depth in the regions marked by the red triangles. The capacity to image optically thick sections of the brain with high resolution and contrast enables the inspection of axonal connections with minimal damage, in contrast with, for instance, microtome sectioning needed for conventional fluorescence microscopy. Airy DL LSM provides attractive opportunities for detailed studies of morphology, neurodegeneration and broader pathogenesis in neuroscience.

**Fig. 6.**
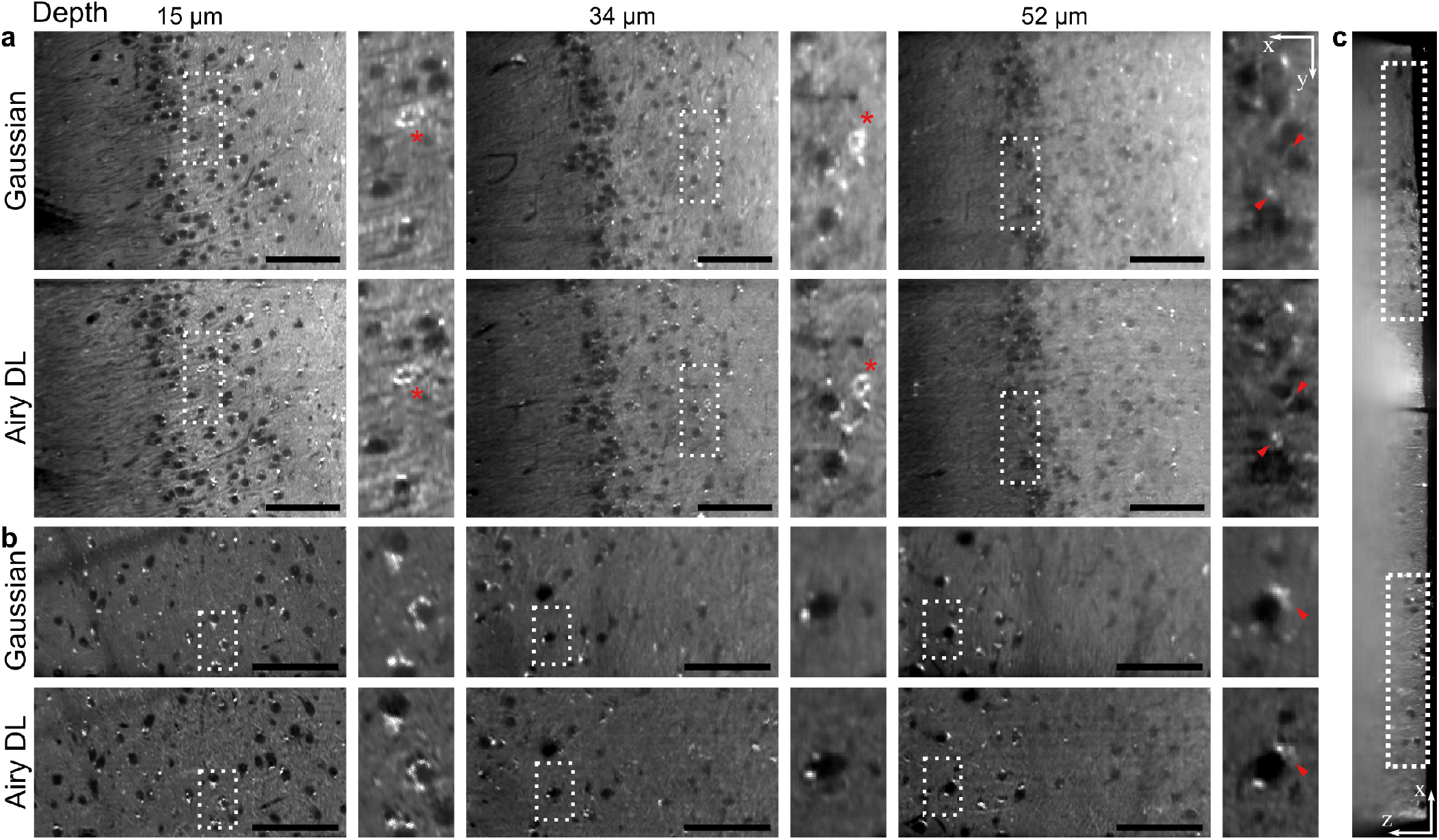
Excised brain tissue from a mouse expressing enhanced YFP in parvalbumin-positive (PV+) neurons. (a, b) Enface sections in depth imaged using a Gaussian and Airy LSM with deep learning deconvolution. Sections correspond to the (a) hippocampus and (b) deeper cortical regions. Insets to the right correspond to the regions marked by a dashed rectangle. Red asterisks mark PV+ neurons. Red triangles mark regions that illustrate enhanced performance from the combination of an Airy beam and deep learning at depth. (c) Cross-sectional image (*z* is depth) showing the regions displayed in (a) and (b). Scale bars are 100 *μ*m.

### Multiphoton LSM with a Bessel beam

We have demonstrated our DL method with Airy LSM because of the facile control of the PSF, enabling Gaussian LSM ground truths. However, our method is also readily adaptable to deconvolution of other PSFs, such as the Bessel beam which is common to many micropscopy methods (6, 8) and underpins lattice LSM methods (9). Towards this, we briefly demonstrate the application of DL to a multiphoton LSM system using a Bessel beam, generated using an axicon lens (11). The setup and data acquisition methods are presented in detail in (41). Figure 7 illustrates the low-resolution Bessel images (LR) deconvolved with a DL network trained on a manually matched Bessel PSF. Figure 7(a) clearly illustrates the capacity to remove the side lobe structures generated by the Bessel beam. The resolution improvement in Fig. 7(a) can be quantified as the FWHM of the PSF profiles. Resolution improved from 2.3 *μ*m in LR to 1.6 *μ*m in DL images, which is consistent with the improvements in the Airy LSM (Fig. 3).

**Fig. 7.**
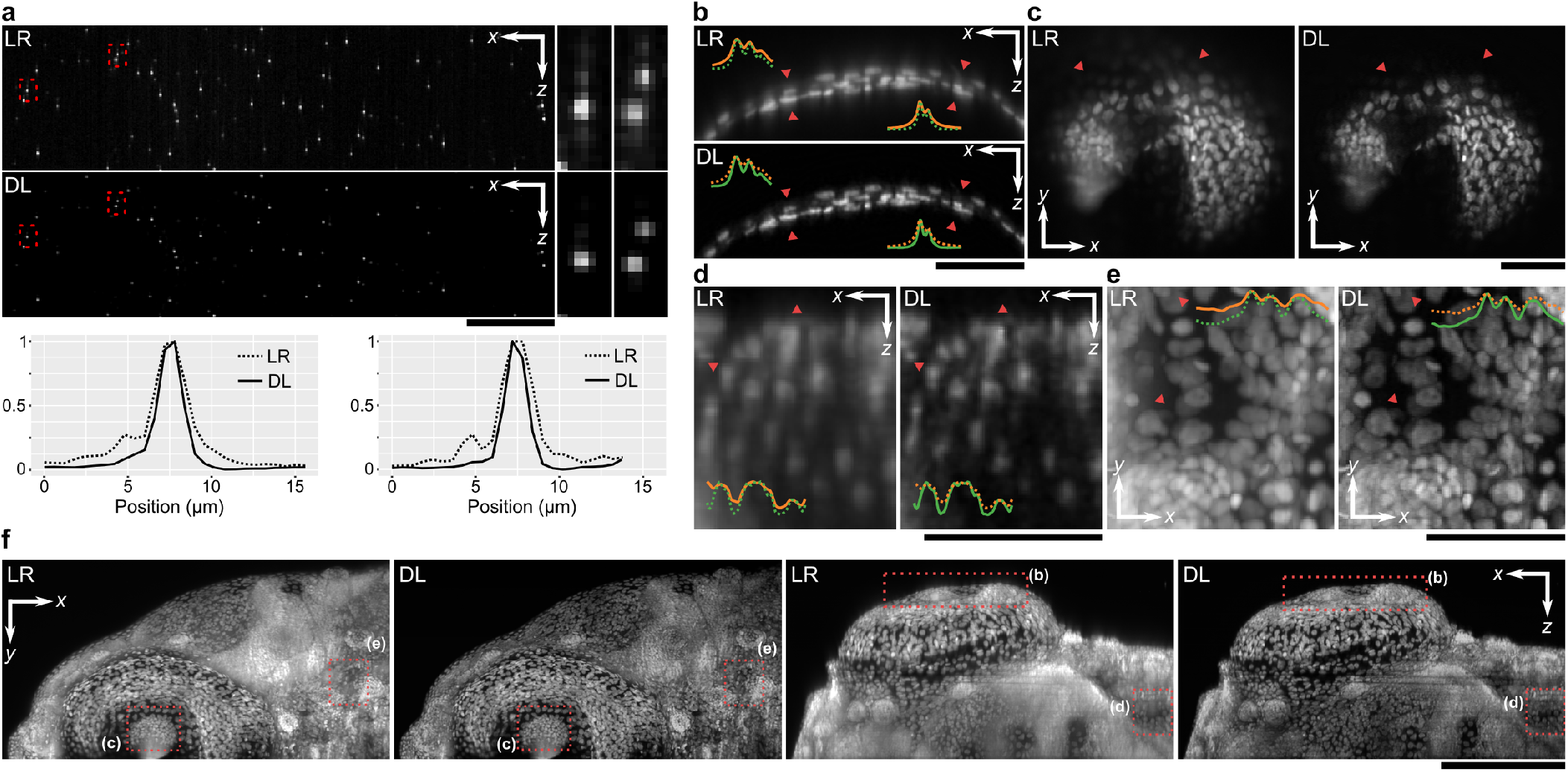
Learned deconvolution in a multiphoton LSM with a Bessel beam. (a) Fluorescent beads encoded by a Bessel PSF (LR) deconvolved using deep learning (DL). Zoomed in regions correspond to areas marked by the red rectangle. Line plots are representative of the axial intensity profile, showing deconvolution of side lobes. (b–f) Zebrafish embryo sections: (b) cross-sections of the iris; (c) en-face sections of the pupil; (d) cross-sections of the body; (e) en-face maximum intensity projections (20 *μ*m) of the body. Line insets correspond to intensity profiles along the regions marked by red triangles. In (c), red triangles illustrate out-of-plane defocus. (f) Maximum intensity projections of the entire sample in the cross-section and en-face planes. Red rectangles mark locations corresponding to (b–e). Scale bars are 50 *μ*m in (a–e) and 200 *μ*m in (f).

Figures 7(b–f) illustrate DL deconvolution in images of a Zebrafish embryo, which is a common model organism for developmental and cardiovascular studies (41). Figures 7(b–e) show the performance of DL in different sections of the Zebrafish corresponding to regions marked in Fig. 7(f). Image intensities were normalised to the background intensity. It is evident that DL images present sharper contrast and importantly a reduction in haze from the side lobe structures generated by the Bessel beam (e.g., Fig. 7(c)) in both the zoomed-in regions and the maximum intensity projections in Fig. 7(f) The spatial resolution was improved from 2.6 *μ*m in LR to 1.9 *μ*m in DL images (Supplementary Note 10).

In Supplementary Note 12, we further demonstrate the broad applicability of our method by deconvolving data made openly available by others. Specifically, timelapses of a developing Medaka larvae imaged using multiphoton LSM with a Bessel beam, provided by Takabezawa *et al.,* (42) as open source. The full timelapse videos are available as Supplementary Movies S1 and S2.

## Discussion

We have demonstrated that experimentally unsupervised deconvolution using physics priors in lieu of experimental ground truths can achieve powerful and generalised deconvolution, with performance exceeding that of conventional deconvolution methods. The improvements of DL we compared in detail to RL deconvolution, including with TV regularisation, which are among the top performing and commonly used algorithms (3) (Supplementary Note 6). Compared to typical end-to-end approaches, our method simplifies the requirements to solely images that would be collected in a standard experiment and, importantly, does not require investment in new imaging hardware and experiments solely for training the network. This physics-informed learning is a marked contrast to data-driven approaches (13–15), which derive their performance, in part, from sample-specific priors, thus, requiring training for each class of sample. The use of physics theory to train neural networks has emerged in recent works (12), including in correcting for scattering (43) and enabling deconvolution of spatially varying PSFs for diffuser-based imaging (44). It is likely that more complex physics simulations, such as those including the simulation of aberration and scattering, will lead to better-performing networks (12). Accurate physics simulations have already demonstrated substantial improvements in computational methods in light-field microscopy (45), phase unwrapping (20) and structured illumination microscopy (21). Further, aberrations from alignment incorporated in a network via Zernike polynomials have improved phase retrieval (46). In future work, more complex priors may also be matched with the capacity for deeper or volumetric network training, or training with multi-step networks to address individual image enhancement tasks, as demonstrated by RCAN (15) and DSP-Net (16) networks. Whilst our method could be readily extended to 3D deconvolution, 3D network training is memory-intensive and requires costly graphics hardware (Supplementary Note 13). In LSM, for instance, the accuracy of 2D deconvolution was only 2– 3% poorer than that of 3D (Supplementary Note 8), which is an acceptable trade-off for accessible and rapid training and inference.

Many image restoration methods rely on enhancing a finite amount of information carried by an image, which is naturally an underdetermined inverse problem. As such, constraints must be placed on inversion to achieve a unique and accurate result (30). For example, regularisation schemes, such as total variation or *L*_2_-norm minimisation, have been used with deconvolution. Deep learning achieves this exceptionally well using feature learning by generating outputs that are most similar to expected images, whilst excluding those solutions that are unlikely to be encountered in microscopy. Specifically, we can consider deep learning to comprise physical and content learning (4, 22). Physical learning encourages consistency with the transformation by the imaging system, for instance, the blur from the finite bandwidth of optical elements or the noise in detectors. Content learning encourages output images to resemble features observed in the ground truths related to, for instance, the type of samples or spectral contrast expected. While physics-informed learning simplifies training and enables broad generalisation by favouring physical learning (4, 12), the addition of content learning though accurate experimental ground truths offers the greatest constraints to optimisation and inversion tasks. Thus, content learning will likely still offer the greatest performance, which is evident in cross-modality restoration (14, 15). Though, care must be taken to ensure that accurate image content is recovered through adequate generalisation for a particular type of sample (17) and, further, broader generalisation leads to poorer performing networks for a particular sample set. The challenge in paired data acquisition further limits the broad uptake of such methods, which may become only feasible for routine imaging of samples with fixed optical instruments, for instance, with commercially available microscopes. We discuss these methods in more detail in Supplementary Note 13. It may be possible to augment data-driven approaches with physics priors to reduce the volume of data required and optimise the balance in physical vs. content learning (12). In future studies, a detailed comparison of simulation-based and data-driven methods would be instrumental in determining the right balance between physical and content learning.

Our DL method is particularly useful for LSM with propagation-invariant Airy and Bessel beams, decoding image content with a high contrast, over an extended field of view and additionally provide super-resolution capacity. Specifically, we show that DL is superior to conventional deconvolution in its ability to achieve a more symmetric deconvolved PSF, a PSNR of 30–33 dB compared to 25–30 dB for RL deconvolution, and a 3-fold improvement in the tolerance to a mismatched PSF. Importantly, we also show that the combined Airy DL method exceeds the performance of a conventional Gaussian LSM, exceeding both the resolution and the contrast, as illustrated by the MTF (Fig. 4(f) and Supplementary Note 10). It is important to note that the use of propagation-invariant beams leads to a reduction in contrast due to the distribution of power into the side lobes (5), establishing a trade-off in resolution, contrast and DOF. Scattering through turbid biological tissues and background fluorescence can lead to a reduction in resolution and contrast (Supplementary Note 10). DL, however, has extended both resolution and contrast compared to the Gaussian LSM, which reduces one of the key downsides of propagation-invariant beams and offers new opportunities for rapid multiscale bioimaging, such as in embryology and neuroscience.

Further, the support of DL by contemporary hardware, such as the GPU, promise ultrafast and even real-time processing with a low barrier-to-entry. Airy and Bessel beams can be generated readily using cheap optical components (5, 10, 11). The combination of DL with compact and inexpensive LSM may provide a powerful and accessible tool for bioimaging. This may further democratise other precision LSM setups, such as the lattice light sheet (9) and more broadly other microscopic approaches that employ deconvolution.

## Conclusion

We have demonstrated learned deconvolution of propagation-invariant beam shapes in LSM, leading to the recovery of high-resolution and high-contrast information over extended fields of view, with superior performance to conventional deconvolution methods. Our method is further distinguished from many other learned methods in that it is experimentally unsupervised and requires no high-resolution ground truths, which has led to good performance and generality. The application to embryology and neuroscience show attractive potential across a range of bioimaging studies. Our method, which is provided to the reader as open source, may enable ready implementation accessible to many imaging setups.

## Materials and Methods

### Network

Let us consider the LSM imaging process described by *y* = *H(x)* + *ε*, where *x* is the ground truth (GT), *y* is the recorded image (LR), *H* is the imaging operator that we seek to invert, and *e* is some noise. The goal of deep learning is to train a network, *G*, such that is provides a good estimate of the ground truth (DL) from LR images, 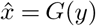. This is achieved by minimising a training loss function (TLF), 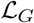, i.e., argmin_*G*_ 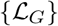. We implement this process using a generative adversarial network (GAN) (23), where *G* is trained concurrently with an adversarial discriminator network, *D*, that seeks to maximise its ability to discriminate between *x* and 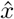. *D* features in 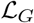, and leads *G* towards optimising the adversarial min-max game (23).

The discriminator, *D*, is trained to minimise the following TLF based on least-squares (47):

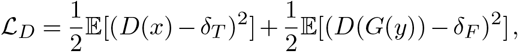

where *δ_T_* and *δ_F_* are the ‘true’ and ‘false’ labels, and 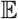 is the expected value evaluated over the entire training batch. We utilise one sided smooth labels, i.e., *δ_T_* = 0.9 and *δ_F_* = 0 to prevent *D* from over-saturating learning from a particular strong feature. *D* is based on the Patch GAN architecture (25) (Supplementary Note 1).

The Generator, *G*, is trained to minimise a linear combination of three TLFs:

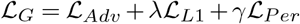

which are the adversarial, *L*_1_-norm and perceptual losses, respectively. Weights *λ* and *γ* are chosen such that 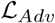 and 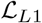 have approximately equal contribution to 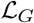 at the end of training (here, *λ* = 30). Perceptual loss acts as a regulariser, and was manually chosen as 1 × 10^-4^ (Supplementary Note 2). *G* is based on a 16-layer residual network (ResNet) (24), inspired by SRGAN (26) (Supplementary Note 1).

We further use multi-objective training (48), i.e., we train two independent discriminators, *D*_1_ and *D*_2_. Using each discriminator, we evaluate a least-squares adversarial loss (47):

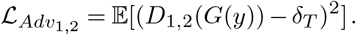

The minimum of the two is used to update the gradients of *G*:

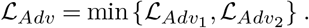

Multi-objective training prevents a singe *D* and *G* being locked into a min-max game over a single strong feature. In such a case, the second discriminator becomes momentarily free of the min-max game, and is able to reestablish a good training gradient.

Pixel loss using *L*_1_-norm establishes spatial consistency between DL (*G*(*y*)) and GT (*x*) images, and is given as:

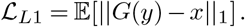

A further perceptual loss is provided with a pre-trained and publicly available VGG-16 (27) network (*V*). Inference of an image with *V* leads to its layers possessing some quantification of image content that is less dependent on its precise pixel values (28). Specifically, we compare LR and DL image content by extracting intermediate features of their *V* inference:

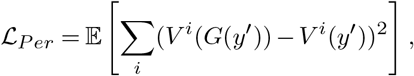

where *i* are the selected layers of *V*.

It is important to note that 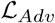 and 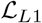 are calculated using the DL and GT images that are simulated based on the known light propagation in the LSM (Fig. 1). Real acquired LR images feature solely in 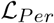. Perceptual loss, in this instance, plays an important role in directing *G* towards good performance in real tissues. If a pixel-wise loss was to be used instead, it would act to make DL images look like LR images, which is counter-productive. Instead, perceptual loss provides two important features (Supplementary Note 2). First, it focuses on preserving content that is present in the LR images, namely, the distribution of spectral density, contrast and brightness, and the low-resolution content that should be unaffected by the deconvolution process. Second, it flags network artefacts, such as fixed pattern noise, with strong increases in loss. This makes it a valuable regularisation parameter to bridge mismatch between the content that can be efficiently simulated and real samples. Perceptual loss has previously been valuable for phase-retrieval in microscopy (49).

### Training and Inference

Training was performed using simulated LR images and their GT, and real LR images of tissue. All inputs and DL outputs were 64×64 pixel grayscale images. LR images were simulated by convolving a sparse or speckle bases (Fig. 1(b)) with a known PSF. The PSF was evaluated by superimposing a Gaussian detection beam orthogonal to an illumination beam. The beam propagation models are described in the subsequent sections. GT images were simulated by convolving the same basis used for LR with a Gaussian spot at 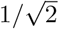 the diffraction limit. Real LR images were retrieved from an LSM volume of a blastocyst in an automated fashion by identifying the brightest region of interest in each cross-section and applying a random spatial shift. Simulated LR and GT images were normalised to a range of (0.1, 0.9) to avoid clipping. The real LR images were scaled to the minimum and maximum intensity of the entire LSM volume.

Training for each beam type was performed using a total of 512 sparse and 512 speckle simulations, and 215 LR images from a blastocyst volume. Of these, a random selection of 32 sparse and 32 speckle LR images were used for validation. Due to the adversarial nature of GAN training, no early stopping was used. This is because adversarial loss is not an indication of convergence. We used a batch size of 8 and trained for 300 epochs. Gradient-based optimisation was performed using the Adam optimiser with learning rates of 1 × 10^-4^ and 4 × 10^-4^ for *G* and *D*, respectively, and hyper-parameters *β*_2_ = 0.5 and *β*_2_ = 0.999. Training was performed using PyTorch and an NVidia RTX 2060, and took approximately 2.5 hours. Inference on widefield images was performed by a 64×64 sliding window with an 8-pixel overlap to avoid boundary issues. Inference took 3 ms for a single LR image, which corresponds to approximately 0.5 s for a widefield LSM image. We replicated training and inference on an NVidia RTX 3090, which reduced the total training time to 1.2 hours, and inference time to 1 ms and 0.2 s for single and widefield sections, respectively. This is comparable to RL deconvolution, which took 0.3 s for a widefield image or 0.5 s with TV regularisation.

### Richardson-Lucy deconvolution

Richardson-Lucy deconvolution was performed in 2D (and 3D in Supplementary Note 8) using a Python interface to ImageJ, which also includes common implementations, such as using total variation regularisation, as part of its Ops framework.

### Imaging system

The imaging setup is illustrated in Figure 1(a). A 488-nm laser source (TOPTICA, Germany) was used for illumination. The beam was expanded with an objective (Obj_B_, RMS10X, 0.25NA, 10X, Olympus) and lens (L1, f:150 mm), and apodised using an aperture. The beam was filtered spectrally using a notch filter (*λ_c_*:480 nm, Δ*λ*:17 nm) and spatially using a pinhole (15-*μ*m diameter) at the objective focus. A half wave plate controlled polarisation to maximise the efficiency of the spatial light modulator (SLM, X10468, Hamamatsu, Japan). Phase and amplitude was controlled by using the SLM in diffraction mode. The desired phase was projected onto the SLM and the desired intensity was modulated by a phase ramp, realising a blazed grating. The first-order diffraction was spatially filtered using an aperture (SF). The beam was relayed by lenses (L2–L5, f:250, 100, 50, 75 mm) and the illumination objective (Obj_I_, 54-10-12, 0.367NA 4X, Navitar). The light sheet was generated by a galvo scanner (Galvo, Thorlabs, NJ) in the Fourier plane. Fluorescence was collected by the detection objective (ObjD, 54-10-12, 0.367NA 4X, Navitar) and relayed to the camera (CAM, Iris 15, Teledyne Photometrics, AZ) by a tube lens (TL, f:200 mm). Fluorescence was filtered using a bandpass filter (BP, *λ_c_*:532 nm, Δ*λ*:50 nm) and a notch filter (NF, *λ_c_*:488 nm).

### Acquisition

LSM volumes were acquired by placing the sample at the focus of the objectives and laterally scanning using a motorised transducer (M-230.10, Physik Instrumente Ltd). Scanning position was synchronised to the camera acquisition, and images were acquired in 0.5-*μ*m steps. Due to the geometry illustrated in the inset in Figure 1(a), lateral scanning was at 45° to the detection plane (also detailed 11 in (38)). Due to this shearing, acquired image stacks were transformed to the physical Cartesian coordinate space using in-house built software (MATLAB, MA) based on interpolation. All LSM images presented in this paper were sampled with an isotropic 0.85 *μ*m pixel size.

### Structured light fields

The description of the beams used in this work is provided in Supplementary Note 14. Briefly, all beam shapes are described in the pupil plane. The Gaussian beam is described by its 1*/e*^2^ waist size (*w*_0_). The Airy beam is described by the 1*/e*^2^ waist size (*w_0_*) of a Gaussian intensity and a scale-invariant *α* parameter that represents the cubic phase modulation rate. The Bessel beam is described by a ring with a 1/*e*^2^ waist size (*w*_0_) and a ring radius (*rr*). Table 1 lists the beam parameters used in this work.

**Table 1.** Illumination beam properties. Res: axial resolution. DOF: depth of focus.

| Beam | Parameters (mm) | Res. ($\mu\text{m}$) | DOF ($\mu\text{m}$) |
| --- | --- | --- | --- |
| Gaussian 1 (G1) | $w_0$: 1.02 | 2.5 | 27 |
| Gaussian 0.5 (G0.5) | $w_0$: 0.510 | 5.3 | 107 |
| Airy 0.5 (A0.5) | $w_0$: 1.38; $\alpha$: 0.5 | 2.6 | 169 |
| Airy 1 (A1) | $w_0$: 1.53; $\alpha$: 1 | 2.5 | 261 |
| Airy 2 (A2) | $w_0$: 1.73; $\alpha$: 2 | 2.5 | 372 |
| Bessel 0.8 (B0.8) | $w_0$: 0.04; $rr$: 0.8 | 3.7 | 510 |
| Bessel 1.1 (B1.1) | $w_0$: 0.055; $rr$: 1.1 | 2.8 | 276 |

### Phantoms

Phantoms were utilised to characterise the system’s PSF and demonstrate DL performance. Phantoms were fabricated from 200-nm diameter green fluorescent microspheres (G200, Duke Scientific, CA) manually mixed with 1.5% agarose. The samples were pipetted into and cured in a 3D-printed sample holder. The sample holder featured a thin fluorinated ethylene propylene (FEP) window for imaging and index matching (RI: 1.344).

### Mouse oocytes and embryos

Female (21-23 days) CBA x C57BL/6 first filial (CBAF1) generation mice were obtained from Laboratory Animal Services (University of Adelaide, Australia) and maintained on a 12 h light: 12 h dark cycle with rodent chow and water provided *ad libitum.* All studies were approved by the University of Adelaide’s Animal Ethics Committee and were conducted in accordance with the Australian Code of Practice for the Care and Use of Animals for Scientific Purposes.

Female mice were administered intraperitoneally (i.p.) with 5 IU of equine chorionic gonadotropin (eCG; Folligon, Braeside, VIC, Australia), followed by 5 IU human chorionic gonadotrophin (hCG, i.p.; Kilsyth, VIC, Australia) 48 h later. Mice were culled by cervical dislocation 14 h post-hCG administration and the oviducts carefully removed. Cumulus oocyte complexes (COCs) were harvested by gently puncturing the ampullae of the oviduct using a 29-gauge insulin syringe with needle (Terumo Australia Pty Ltd, Australia). These COCs were either fixed immediately in 4% paraformaldehye (PFA) or co-incubated with sperm for IVF. In vitro fertilisation (IVF) occurred through co-incubation of matured COCs with sperm in Research Fertilisation medium (ARTLab Solutions, Australia) for 4 h at *37°C*. Resulting presumptive zygotes were transferred into Research Cleave medium (ARTLab Solutions, Australia) under paraffin oil and allowed to develop to the blastocyst-stage (96 h post-IVF)at 37°C in 5%*O_2_*.

Cumulus oocyte complexes (COCs) and blastocyst-stage embryos were fixed in 4% PFA for 30 mins at room temperature, followed by washes in PBV (phosphate buffer saline; PBS containing 0.3 mg/ml of polyvinyl alcohol; PVA). Fixed COCs and blastocyst-stage embryos were mounted under oil on a 3D-printed sample holder with an FEP window for imaging.

### Mouse brain tissue

All animal experiments in Figure 6 were performed in accordance with the United Kingdom Animals (Scientific Procedures) Act of 1986 Home Office regulates and approved by the Home Office (PPL70/8883). Detailed procedures are described elsewhere (50). Briefly, a female adult (8.4 months old) PV-Cre::Ai32 mouse (PV-Cre, JAX008069; Ai32, JAX012569) was used. This mouse expressed channelrhodopsin-2 (ChR2) tagged with enhanced YFP (EYFP) in parvalbumin-positive (PV+) neurons. The mouse was deeply anaesthetised with a mixture of pentobarbital and lidocaine, and perfused transcardially with physiological saline followed by 4% paraformaldehyde/0.1 M phosphate buffer, pH 7.4. After an overnight post-fixation in the same fixative, the brain was immersed in 30% sucrose in phosphate buffered saline (PBS) at 4°C for cryoprotection. The brain was cut into 1-mm thick coronal sections and kept in PBS until imaged. The imaging was focused on the cortex and hippocampus regions.

### Multiphoton LSM

The data and setup used for multiphoton LSM experiments on the zebrafish were used with permission from our previous work (41).

## Acknowledgements

We would like to acknowledge Federico Gasparoli for early support in constructing the imaging system and providing multiphoton data, and Mirna Merkler for preparing the excised mouse brain section. We acknowledge Erik Linder-Norén’s implementations of GANs in Py-Torch (https://github.com/eriklindernoren/PyTorch-GAN) that instantiated the code developed in this project.

## Data Availability

Code used to simulate light-sheet microscopy and perform deep learning is available publicly at https://github.com/philipwijesinghe/learned-deconvolution. All data underpinning this study is available at https://doi.org/10.17630/bf92bc18-0b81-41f7-bd44-d74040af7cf0.

## Author Contribution

PW and KD initiated the project. PW designed the network and imaging theory. SC built the setup and performed imaging. PW performed image processing, network training and image analysis. DJXC and KRD prepared the embryo samples, and provided training and embryo image analysis. SS provided brain tissue, training and analysed brain data. PW wrote the manuscript with input from all authors. All authors reviewed and approved the work. KD supervised the project.

