## Supplementary Notes for "Experimentally unsupervised deconvolution for light-sheet microscopy with propagation-invariant beams"

### Supplementary Data: Learned deconvolution using physics priors for light-sheet microscopy with propagation-invariant beams.

**Philip Wijesinghe[1], Stella Corsetti[1], Darren J.X. Chow[2,3,4], Shuzo Sakata[5], Kylie R. Dunning[2,3,4], and Kishan Dholakia[1,6,7]**

[1]SUPA, School of Physics and Astronomy, University of St Andrews, North Haugh, St Andrews, Fife, KY16 9SS, UK
[2]Robinson Research Institute, Adelaide Medical School, The University of Adelaide, Adelaide, SA, Australia
[3]Australian Research Council Centre of Excellence for Nanoscale BioPhotonics, The University of Adelaide, Adelaide, SA, Australia
[4]Institute for Photonics and Advanced Sensing, The University of Adelaide, Adelaide, SA, Australia
[5]Strathclyde Institute of Pharmacy and Biomedical Sciences, University of Strathclyde, Glasgow, G4 0RE, UK
[6]School of Biological Sciences, The University of Adelaide, Adelaide, South Australia, Australia
[7]Department of Physics, College of Science, Yonsei University, Seoul 03722, South Korea

#### Supplementary Notes:

**1 Network architecture** **2**

**2 The role of the saliency constraint using perceptual loss** **3**

**3 Is a generative adversarial network required?** **4**

**4 Network architecture: ResNet vs. U-Net** **5**

**5 Deconvolution in simulated beads** **6**

**6 Comparison of deconvolution methods** **7**

**7 Effect of noise and background fluorescence on deconvolution** **8**

**8 Validity of two-dimensional deconvolution** **9**

**9 Accuracy of fluorescence quantification with deep learning** **10**

**10 Quantification of resolution in biological images** **11**

**11 Widefield imaging of embryos** **12**

**12 Application of learned deconvolution to open-source data: Bessel-beam LSM** **13**

**13 Comparison of learned deconvolution with published networks** **14**

**14 Structured light fields** **17**

**15 Supplementary Movies** **19**

#### Supplementary Note 1: Network architecture

Figure S1 illustrates the architecture of the Generator and Discriminator networks. The Generator architecture is inspired by SRGAN (1). Specifically, we use a network architecture based on residual learning (ResNet) (2). Greater network depths are beneficial to many computational tasks, including image transformation. However, depth is associated with difficulties in training due to vanishing gradients as the loss is back-propagated across many layers (1, 3). ResNet is an implementation that achieves high network depths with stable gradients by implementing skip connections in each residual block. Skip connections provide identity mapping throughout the network, without the network itself having to learn these mappings across its many convolution filters (4). The network comprises an initial convolution layer that generates 64 features in the image tensor. Then, we use 16 residual blocks that comprise two convolution layers followed by batch-normalisation, and a Parametric Rectified Linear Unit (PReLU) activation function. The convolutional layers support 64 feature maps and comprise small filters. Skip connections are implemented between each residual block and, further, between the initial and before the final convolutional layer. The network embodiment offers the benefits associated with a high network depth, whilst assuring stable training.

The Discriminator is based on a PatchGAN architecture (5). Briefly, the input DL image and the ground truth are concatenated as two features of the same image tensor. Following this, several layers, comprising a convolutional layer followed by a Leaky Rectified Linear Unit (LeakyReLU) activation function, reduce the spatial dimension of the images and expand the feature space. The network terminates with a 4×4 image with 512 features that are combined to form a 4×4 label image. The network has local receptive fields, such that local areas between the DL and GT images are compared. This leads to the Discriminator focussing on higher frequency image content (5), which is of benefit to super-resolution tasks.

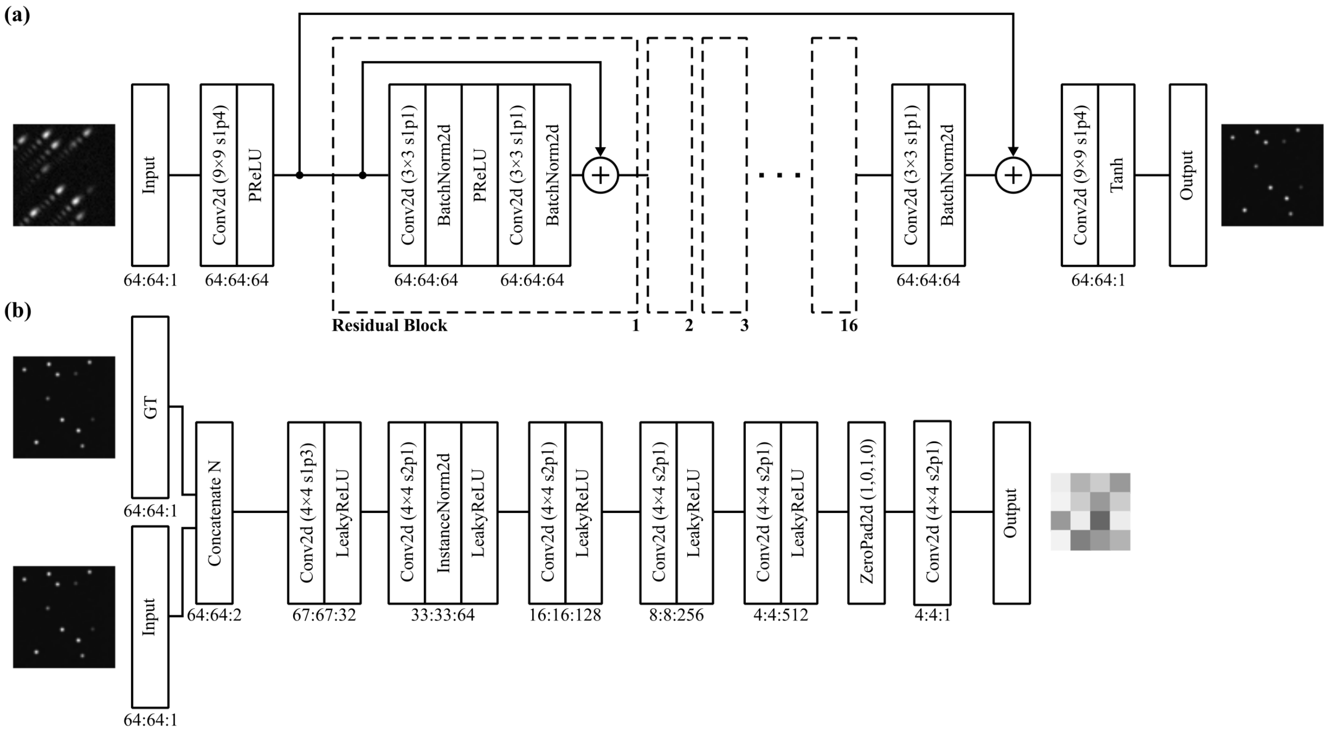


**Fig. S1.** (a) Generator. (b) Discriminator. Values below each block represent the horizontal, vertical and feature dimensions of the output tensors, respectively.

#### Supplementary Note 2: The role of the saliency constraint using perceptual loss

Figure S2 illustrates deep learning deconvolution with various levels of saliency constraint via a perceptual loss between real LR and DL images. The DL images are compared to an image taken in the same region with a Gaussian LSM for reference. When a saliency constraint is not used ($\gamma = 0$), the network fails to learn well the intensity distribution content expected in the image. Specifically, we can see that regions with a low intensity (for instance, that marked by the red asterisk) are disproportionately attenuated by the network. This is likely because they fail to reach some threshold in activation of network layers. Often, only the bright regions are preserved. When a relatively high weight is given to saliency ($\gamma = 1 \times 10^{-2}$), we can see that features start to present blurring in the direction of the Airy side lobes (top right to bottom left). This leads to a loss of resolution, which is expected as the network tries to optimise DL images to match that of the LR input. However, a balanced choice of saliency weight, here $\gamma = 1 \times 10^{-4}$, leads to a preservation of image intensity, even in the low-intensity regions, and at the same time the effective learning of resolution. Control of the intensity distribution in the spatio-spectral domain is important to accurate network training and inference, and has recently received attention in computational microscopy (6–8).

We have further observed that perceptual loss is able to efficiently recognise network artefacts, such as those along the image boundaries, or the fixed pattern noise generated by the finite size of the convolution kernels. As such, we believe it to be useful in network training to discourage the generation and amplification of artefacts, which are likely to be picked up by the discriminator and may lead to poor gradients from adversarial loss.

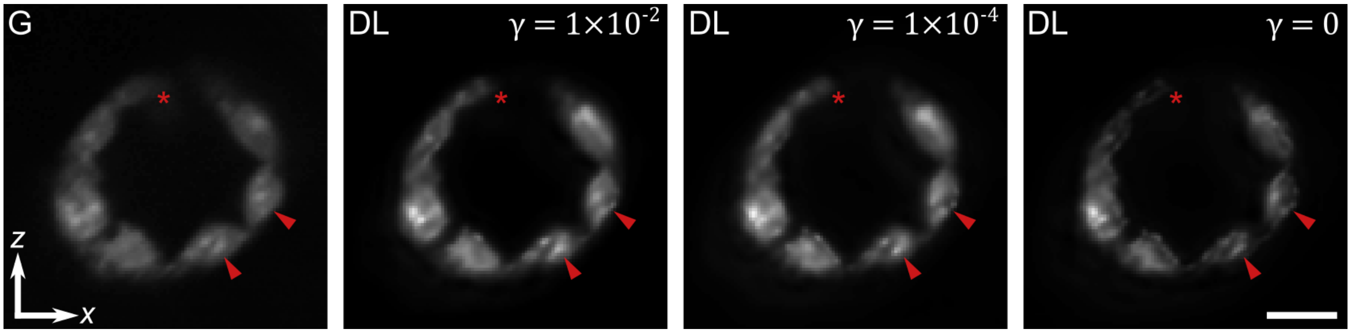


**Fig. S2.** LSM images of a blastocyst. Taken with a (G) Gaussian beam, and Airy beam with deep learning (DL) deconvolution with perceptual weights $\gamma$ of $1 \times 10^{-2}$, $1 \times 10^{-4}$, and $0$. Scale bar is 20 $\mu$m.

#### Supplementary Note 3: Is a generative adversarial network required?

Generative adversarial networks have received much attention in image transformation tasks (1, 5), including in microscopy (9, 10). This is because adversarial networks can learn and emphasise similarity metrics that are not feasible to capture using pixel-wise loss functions, such as the mean-squared error or the $L_1$ norm. In Figure S3, we compare the performance of our network with and without adversarial loss. Specifically, we have trained an additional network using the same training data, however, without training the Discriminator and, subsequently, removing its loss function from $\mathcal{L}_G$. In Figure S3, we can see that when compared to the Gaussian image, a GAN-trained network is able to resolve fluorescent featured with greater visual detail. Further, in the inset in the right-hand image, we can see an improved demarcation between individual cells. This visual improvement in high spatial frequency features is consistent with previous observations (1, 11).

GAN networks can be challenging to train because of unstable and sometimes vanishing TLF gradients (3, 12). We have observed that a balance between the adversarial and $L_1$-norm TLF values, and the addition of dual-discriminator training, leads to consistently stable training. Further, the removal of Discriminator training and loss calculation results in approximately a 5% improvement in training speed. This is because the majority of computational time is dedicated to the transmission of image data between the GPU memory and disk.

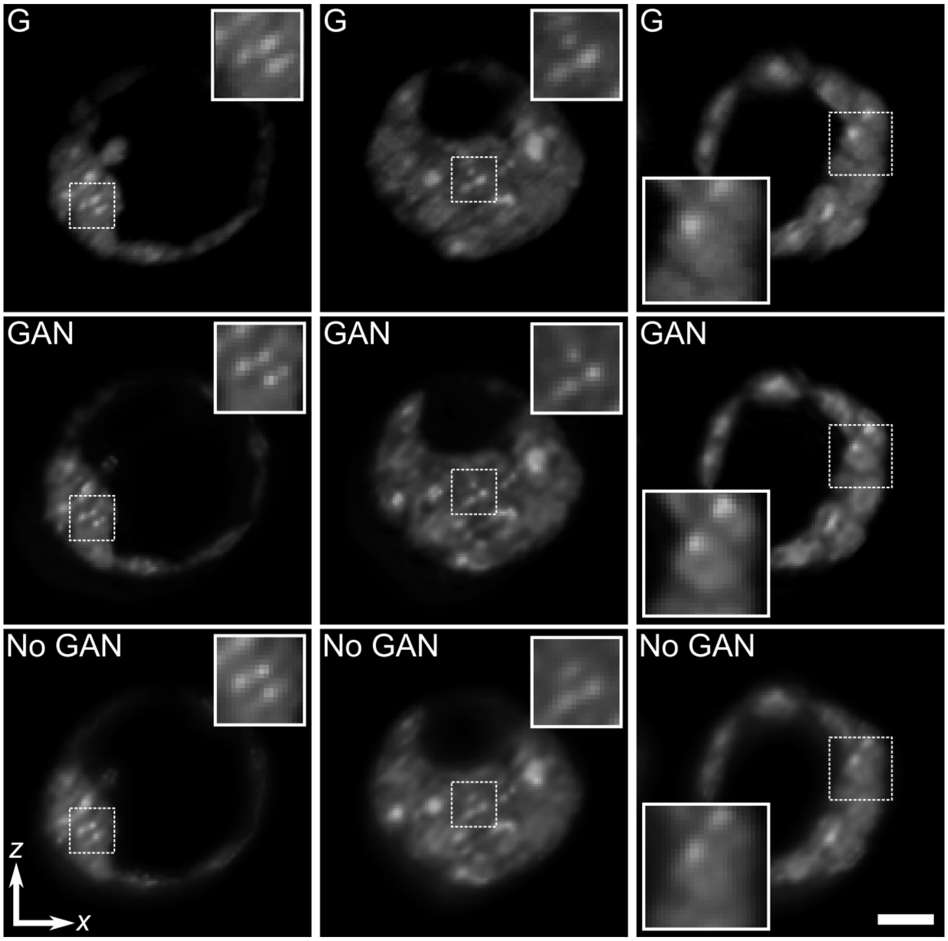


**Fig. S3.** The contribution of generative adversarial network (GAN) training to deconvolution image quality in mouse blastocysts. (G) Gaussian LSM image. Airy LSM image deconvolved using (GAN) adversarial, and (No GAN) adversarial-free networks. Scale bar is 20 $\mu$m.

#### Supplementary Note 4: Network architecture: ResNet vs. U-Net

The selection of network architecture is important to the optimisation of image transformation tasks. Two classes of networks have been prominent in microscopy literature (13). Specifically, the U-Net (or a variation of an encoder-decoder architecture), for instance in (9, 14), and ResNet (or a variation of residual block architectures), for instance in (15, 16). Typically, U-Nets exceed in tasks that require image transformation where the input and output may have substantially different features or styles, for instance in cross-modality transformations (9). ResNets, however, tend to preserve spatial consistency between images because of their smaller receptive fields (1).

In Figure S4, we visually compare the performance of two common implementations of a U-Net (based on pix2pix (5)) and ResNet (SRGAN (1)) as the Generator of our deep learning method. Specifically, we learn deconvolution of an Airy LSM image with two networks using identical hyperparameters and training data. It is clear that the ResNet outputs are more consistent with the reference Gaussian LSM image in the relative intensity distribution, fluorescent feature shape and the image resolution. The U-Net outputs present visibly sharper high-intensity features, which may be suggestive of a higher image resolution. However, there is a focus on high-intensity features and a suppression of low-intensity areas. Interestingly, these visual features seem to be consistent with the outputs of pre-trained U-Net networks in Wang et al. (9) (CMSR, Figure 2). In this work, we have chosen to use the ResNet architecture to emphasise consistency with the reference Gaussian images, which is promising towards quantitative image analysis, evidenced in Figs. 5 and S9.

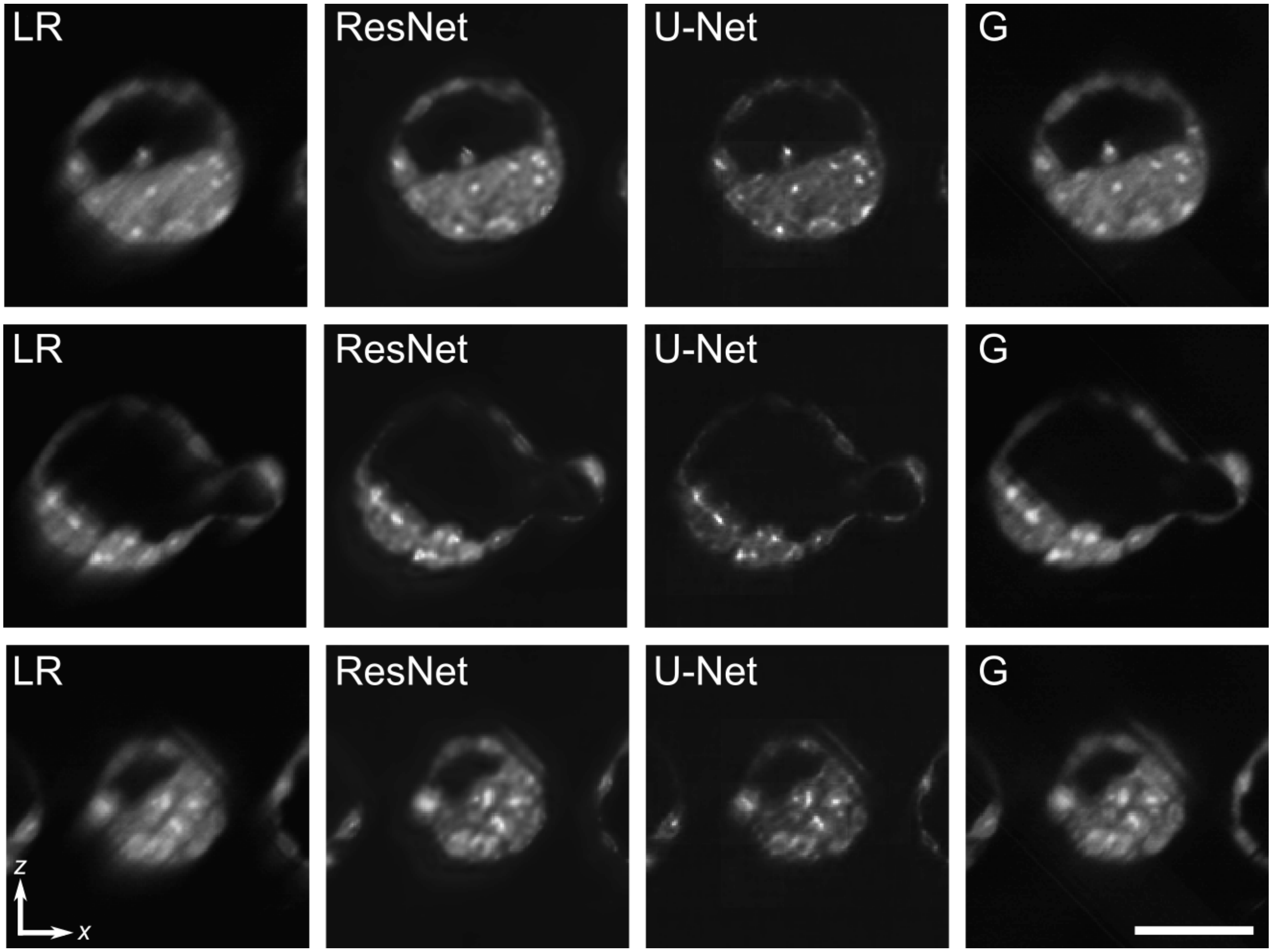


**Fig. S4.** Comparison of ResNet vs U-Net Generator architectures in learned deconvolution of Airy LSM images of mouse blastocysts. (LR) Low-resolution Airy LSM images and (G) reference Gaussian LSM images. Scale bar is 50 $\mu$m.

#### Supplementary Note 5: Deconvolution in simulated beads

Figure S5 visualises the performance of RL, TV and DL deconvolution methods on simulated beads images with various beam shapes (supplement to Fig. 3). We can see that the outputs of RL and TV deconvolution remove the structure of PSF side lobes, however, some elevated intensity remains. This is in contrast to DL, which visually enhances the deconvolved bead structure and matches more closely to the GT.

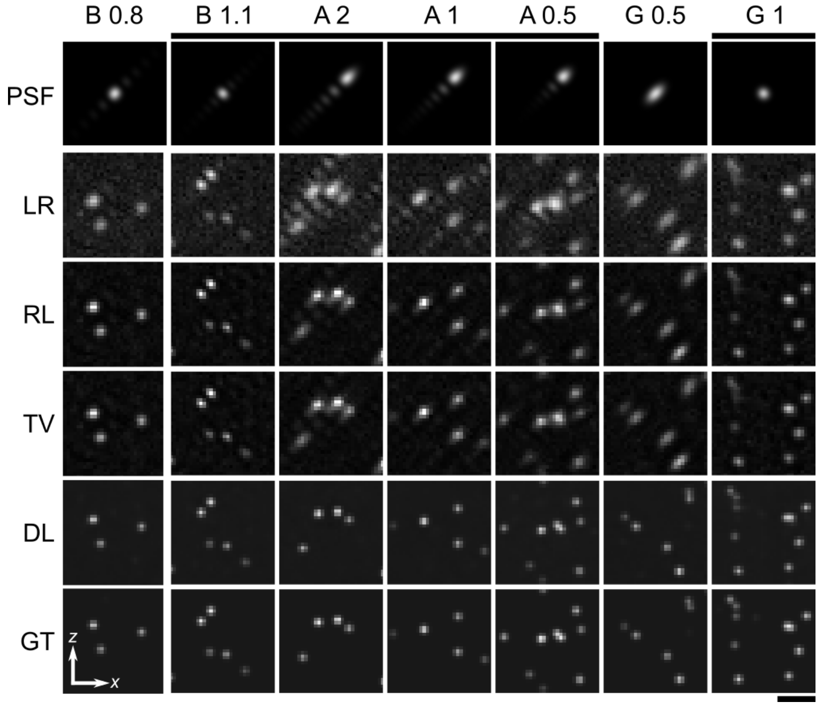


**Fig. S5.** Comparison of deep learning (DL), Richardson-Lucy (RL) and Richardson-Lucy with total variation regularisation (TV) deconvolution in simulated bead data. Deconvolution of low-resolution noisy images (LR) simulated using beams (B: Bessel; A: Airy; G: Gaussian) visualised by their point-spread function (PSF) (parameters listed in Methods) using RL, TV and DL algorithms and compared to the ground truth (GT). Scale bar is 10 $\mu$m.

#### Supplementary Note 6: Comparison of deconvolution methods

Figure S6 compares our deep learning deconvolution method to conventional direct and iterative methods featured in the 'DeconvolutionLab2' open-source package (17). Figure S6 illustrates methods, including: a direct Tikhonov Regularized Inverse Filter (TRIF); Richardson-Lucy deconvolution (RL); RL deconvolution with Total-Variation regularisation (TV); Non-negative Least Squares method (NNLS); Iterative Constraint Tikhonov-Miller (ICTM) and our deep learning method (DL). The input images (LR) comprise ground-truth (GT) images of mouse embryos (421 images) synthetically blurred using the Airy ($\alpha = 1$) beam with 2% additive Gaussian and Poisson noise. The performance of these methods in scattering biological images is similar to that observed previously (17). Specifically, direct inversion (TRIF) performs poorly in low-SNR regions, while iterative regularised schemes perform within a range of a few dB based on the appropriateness of the edge- or smoothness-preserving regulariser. The DL method is superior to these conventional methods, with more than two-fold improvement in the PSNR (>3 dB) to the leading method.

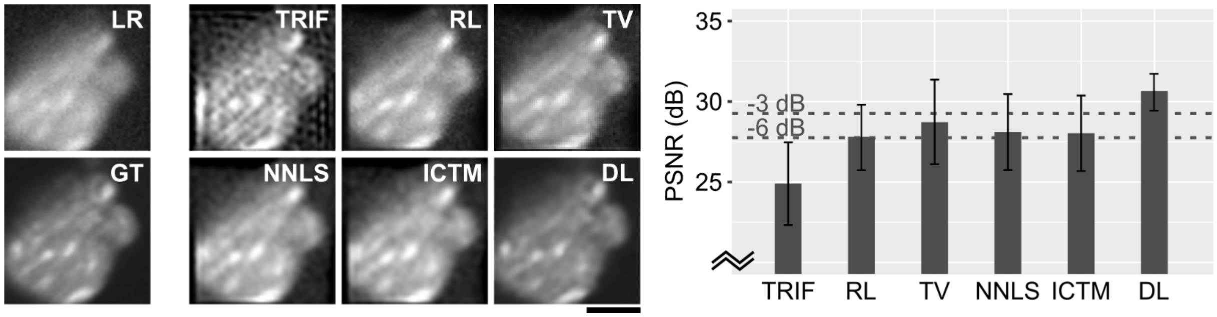


**Fig. S6.** Comparison of deep learning with conventional deconvolution methods. (left) Representative, simulated low-resolution Airy ($\alpha = 1$) LSM image of an embryo, deconvolved using: a direct Tikhonov Regularized Inverse Filter (TRIF); Richardson-Lucy deconvolution (RL); RL deconvolution with Total-Variation regularisation (TV); Non-negative Least Squares method (NNLS); Iterative Constraint Tikhonov-Miller (ICTM) and our deep learning method (DL). (right) Mean peak signal-to-noise ratio (PSNR) values for each method. Dotted lines show -3dB and -6dB reduction in PSNR compared to DL, corresponding to 2$\times$ and 4$\times$ poorer SNR, respectively. Scale bar is 20 $\mu$m.

#### Supplementary Note 7: Effect of noise and background fluorescence on deconvolution

Figure S7 quantifies the effect of noise and background fluorescence on the performance of deep learning deconvolution. We utilised 140 high-resolution Gaussian LSM regions of interest as a ground truth (GT) for simulation. First, we investigate the effect of additive Gaussian and Poisson shot-noise on the reconstruction quality. Low-resolution (LR) images were generated by convolving the GT with the Airy 1 PSF and adding noise proportional to 0–20% of the image intensity. The images were processed with RL and DL deconvolution and quantified via PSNR with respect to the GT. Figures S7(a) and (c) illustrates representative images and the quantified PSNR, respectively. It is clear, visually and quantitatively, that DL reconstructs images closer to the GT. This improvement is consistent with increasing noise.

We also investigate the influence of background fluorescence on reconstruction quality. Specifically, it is signal from out-of-plane of recording that is not modified by the expected PSF of detection. LR images were generated similarly to previous results, by convolving the GT with the Airy 1 PSF. Then a random low-resolution background signal was generated by convolving a random array with a 40 $\mu$m Gaussian kernel. This was added to the LR image, notably without convolution with an Airy PSF, with a proportional contribution in the range of 0–50% of the total image intensity. Finally, a constant 5% Gaussian and Poisson noise was introduced. Figures S7(b) and (d) illustrates representative images and the quantified PSNR, respectively. The elevated background intensity is evident, obscuring the image content. Similarly, DL reconstructs images closer to the GT, and consistently with increasing background.

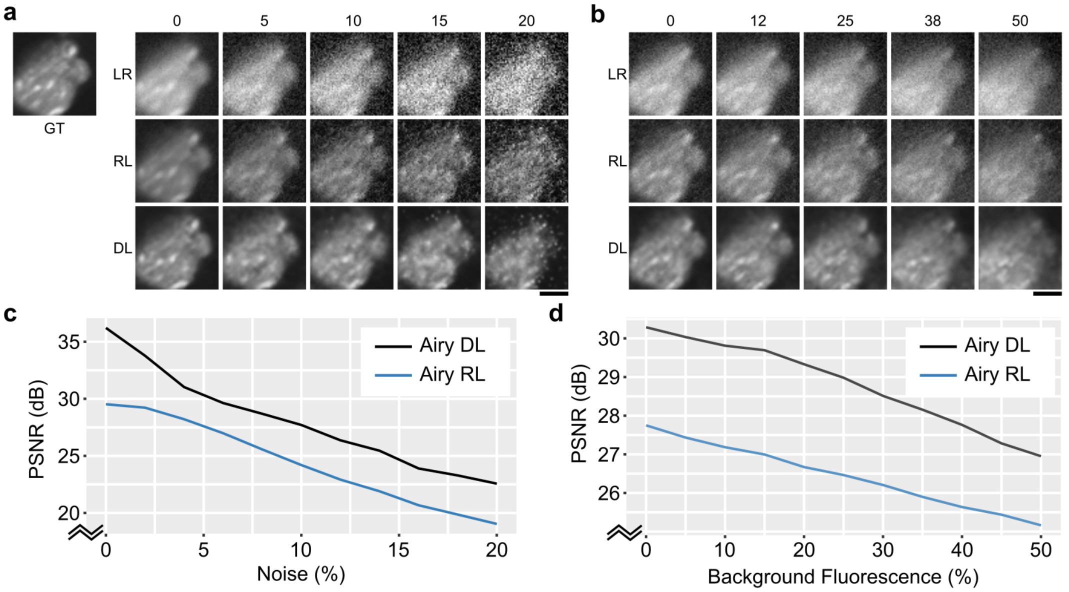


**Fig. S7.** Effect of noise and background fluorescence on deconvolution. (a) Images reconstructed with increasing Gaussian and Poisson noise, and (b) background fluorescence. (c) and (d) PSNR corresponding to images in (a) and (b). Scale bars are 20 $\mu$m.

#### Supplementary Note 8: Validity of two-dimensional deconvolution

Microscopy imaging is a three-dimensional (3D) process. Therefore, it is pertinent to insist that deconvolution and subsequent image analysis should take into account the 3D structure of the image and the 3D system response. However, the added dimensionality over a two-dimensional (2D) problem introduces a substantial computational burden. Therefore, 2D image reconstruction is still prominent in volumetric imaging and when evaluating new reconstruction methods (9, 10, 14). In most microscopy methods, resolution is highest transverse and lowest normal to the axis of the detection. For instance, resolution can be several-fold higher out-of-plane of the camera image than in-plane in epifluorescent setups. This is also true for LSM in its axis of detection.

The errors that arise from using 2D deconvolution of a 3D PSF are due to two main factors. First, deconvolution assumes that the 2D PSF is invariant with defocus, *i.e.,* a cross-section of a 3D PSF defocussed from its centre is equivalent to an intensity-scaled 2D PSF. This means that the contribution of a scatterer that is defocussed from the plane of deconvolution is equivalent to that of a lower-intensity scatterer in focus. Second, no enhancement in resolution is thus possible in the axis normal to that of deconvolution.

To minimise these errors, we have performed deconvolution in the cross-sectional plane that includes the detection axis of LSM. Specifically, 2D images that we deconvolve include the Airy PSF side lobes that are co-axial to the detection, and one of the transverse coordinates (corresponding to the plane of the inset in Fig. 1). This ensures that the resolution is highest out-of-plane (here, the $y$ coordinate), and the discrepancy between a 2D and 3D PSF is minimal. This is evident in Fig. S8(a), showing cross-sections of a 3D PSF and an equivalently scaled 2D PSF, with respect to the defocus of the PSF in the $y$ coordinate. The error is visualised in Fig. S8(a), and illustrates the correspondence of the PSFs. The total error over the 3D extent of the PSF is 2.3% ($\sqrt{\mathrm{MSE}}$).

Figure S8(b) further illustrates the outputs of 2D and 3D deconvolution using the RL method on Airy LSM experimental images of beads. The error map shows areas where the intensity of a 2D-deconvolved image under- and over-estimates that of the 3D-deconvolved image. This is expected because 3D deconvolution enhances resolution out-of-plane (here, in $y$), and the difference between a low- and high-resolution image has a characteristic low-high-low profile, which can be seen in the $xy$-plane insets in Fig. S8(b). The total mean error in deconvolution was approximately 2–3% ($\sqrt{\mathrm{MSE}}$) when thresholding ($> 10\%$) areas of intensity that correspond to the beads, and much lower otherwise.

These results illustrate that 3D deconvolution is preferred in volumetric imaging, however, 2D deconvolution is acceptable with a minimal error. Our method is readily extendable to 3D due to the multidimensional nature of tensor algebra in deep learning. However, it is burdened in requiring greater GPU memory. We have designed our network to operate within 4–6 GB of RAM available to most affordable consumer GPUs, with training taking approximately 1–2.5 hours. Similar residual block architectures have been demonstrated in 3D (15), however, requiring up to 12 GB RAM for training and approximately 12-h training time for similar number of training image pairs. Inference times were also $\sim 0.5$ s here, and approximately 4–5 s for similar sized images in 3D, in (15). However, with the remarkable recent advances in consumer GPUs and cloud processing capacity, the translation to 3D deconvolution using deep learning is a necessary and soon-to-be accessible step.

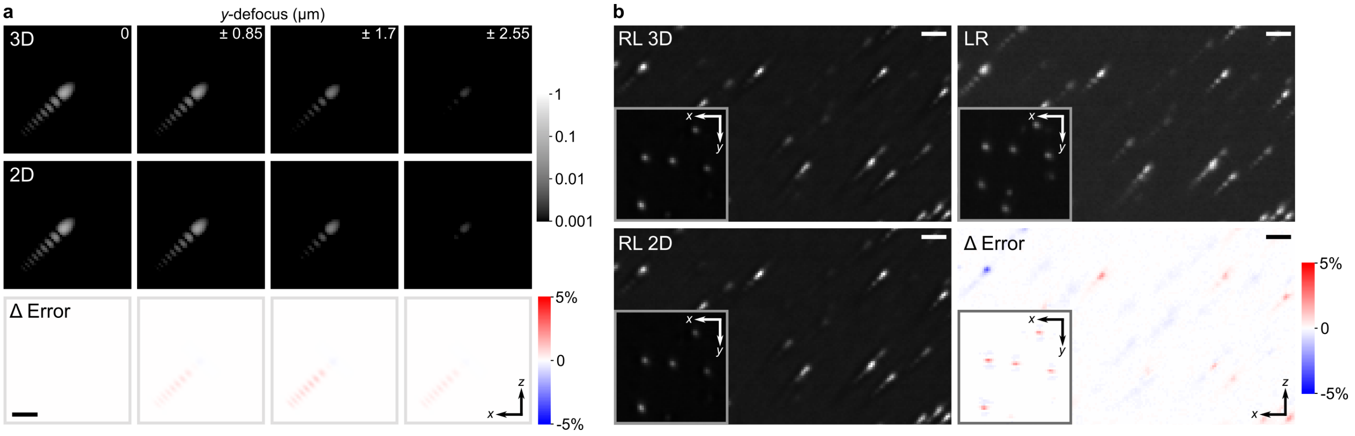


**Fig. S8.** Comparison of 2D and 3D deconvolution of Airy LSM volumes of beads. (a) Comparison of cross-sections of a 3D Airy PSF with a 2D PSF with a scaled intensity, with respect to a defocus in $y$. (b) Richardson-Lucy deconvolution in 3D (RL3D) and 2D (RL2D) from an Airy LSM image (LR) and their relative error. The insets are $xy$ sections normal to the $xz$ cross-sections. Scale bar is 10 $\mu$m.

#### Supplementary Note 9: Accuracy of fluorescence quantification with deep learning

Figure S9 evaluates the capacity of RL and DL methods to recover accurate fluorescent intensity distributions from Airy LSM images of mouse blastocysts. The structured and asymmetric PSF produced by the Airy beam encodes the fluorescence in the recorded images such that local intensities no longer correspond accurately to the surrounding fluorescence signal. This is in contrast to Gaussian illumination schemes, where intensity is monotonic with distance from the fluorescing source. Specifically, because of the oscillating high- and low-intensity areas of the PSF, local pixels may rapidly vary in intensity based on the precise distribution of fluorescent signal and the sectioning. Therefore, deconvolution is necessary to recover a spatially accurate fluorescence distribution. Figure S9 illustrates the performance of RL and DL deconvolution compared with Gaussian LSM reference images. The line plots on the right-hand-side quantify the distribution in the corresponding images. It is evident that DL outputs are more consistent with the Gaussian LSM profiles. In the top row, the DL profile follows closely to G, demonstrating a high contrast between two brightly fluorescent features. The RL profile follows closer that of the LR image with minor enhancement. In the middle row, the DL profile is capable of picking out an additional feature to the left of the largest peak, which is consistent with G. In the bottom row, the DL output similarly shows increased contrast consistent with G. Some variations in the profile are still seen in G. It is important to note that the Gaussian LSM images are not a perfect ground truth because of the much lower DOF and potential movements in the embryos during imaging, which can be seen in the full 3D volumes available under Data Availability.

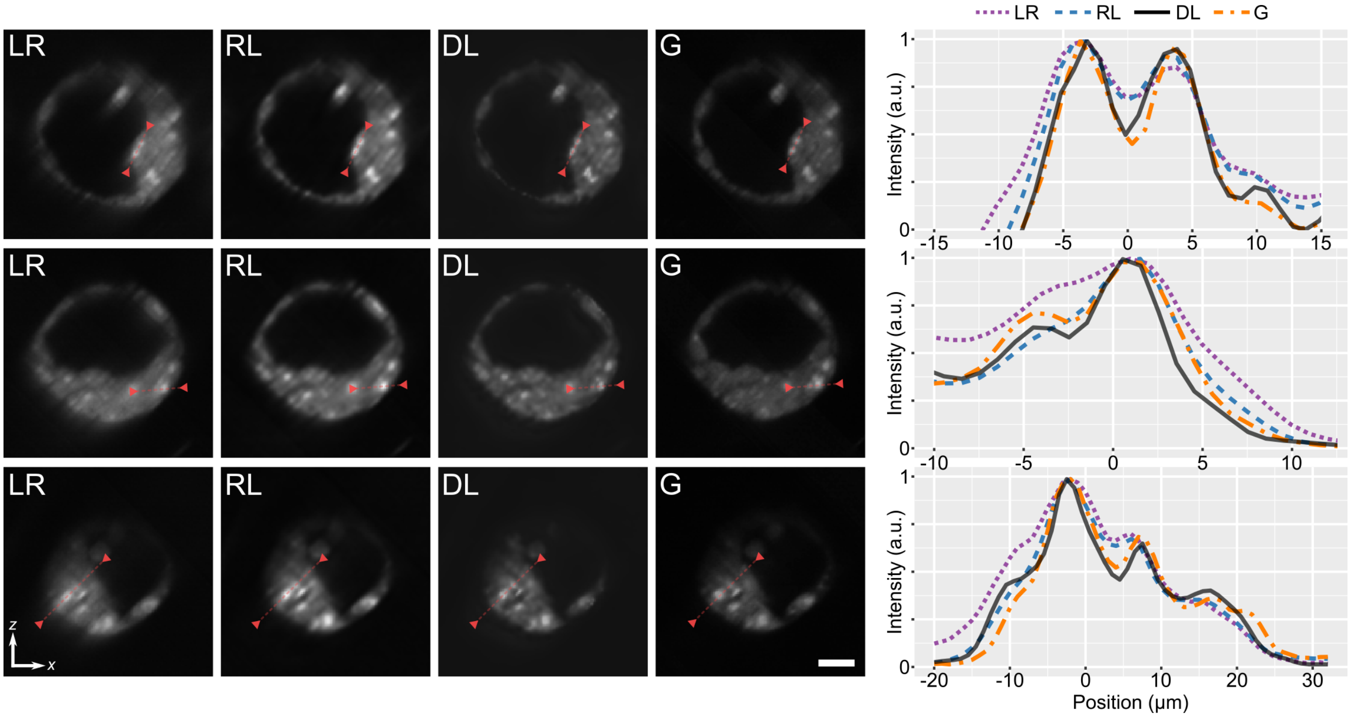


**Fig. S9.** Capacity of deep learning (DL) and Richardson-Lucy (RL) deconvolution in recovering accurate fluorescence intensity from Airy LSM (LR) images compared to Gaussian LSM (G) reference. (left) $xz$ cross-sectional images of mouse blastocysts. The red arrows and line mark the (right) corresponding line profiles. Scale bar is 20 $\mu$m.

#### Supplementary Note 10: Quantification of resolution in biological images

In this work, we quantify resolution in biological tissues using Fourier analysis. When analysing images of beads (Fig. 4), resolution can be quantified readily via the spatial Fourier transform of the image intensity. This is because the intensity signal comprises the convolution of the system PSF with the sub-resolution scatterers that approximate impulse functions, and the signal-to-noise ratio is sufficiently high. In this instance, the Fourier transform sufficiently approximates that of the PSF alone, and relates to the optical transfer function of the imaging system. In biological images, however, the signal comprises an intensity distribution of the sample with its own broad power spectral density convolved with the system PSF, with often additional noise because of the lower fluorescent efficiency of biological labels. Thus, the contribution of the sample and noise must be attenuated to enable Fourier analysis of the PSF alone. We do so in several steps. First, we crop the images to a region or volume of interest that correspond to the sample at focus and sufficient sample contrast. Figure S10(a) illustrates an example of the selected regions. Biological images naturally present higher low-spatial-frequency content (13). We remove this content using a high-pass filter with a cut-off at 0.1 (1/$\mu$m) or a period of 10 $\mu$m for all images. This was determined visually to remove substantial sample influence whilst exceeding the expected system resolution. The high-pass filtered images are shown in Fig. S10(a). Second, the noise, which presents often as a uniform increase across all spatial frequencies, is normalised. Specifically, we select a region in the images with no biological signal. Then, we attenuate its high-frequency components through a low-pass filter with a cut-off determined as the value that reduces the noise power-spectral density to that below 5% of the total for the whole image. Whilst this value is determined individually for each image, all cut-off values were below a period of 0.6 $\mu$m and, thus, were assumed to not greatly degrade the PSF. Figure S10(a) shows an example of the noise regions outlined in red.

The filtered images (such as in Fig. S10(a)) are then Fourier transformed along the dimension that corresponds to the directions of the PSF side-lobe structures. We have averaged the Fourier spectra from 40 images from different regions for each image and sample type to reduce noise. The magnitude of the Fourier transforms, or the estimate of the modulation transfer function (MTF), for each image and sample type are presented in Figs. S10(b–e). The shapes of MTFs correspond to those expected from theory, presented in (Fig. 4). We mark the 5% cross-over value for each MTF, which corresponds approximately to the FWHM of the PSF. We note that the apparent resolution in biological samples is lower than that set by the imaging system and our observations in beads. This is because of several reasons. First, the contribution of the sample is challenging to remove in its entirety without affecting the PSF. Second, the scattering and aberrations through turbid tissues leads to a broadening of the system transfer function and the reduction in resolution. This is especially evident in the brain. The specific plots selected correspond to the images and data presented in the manuscript.

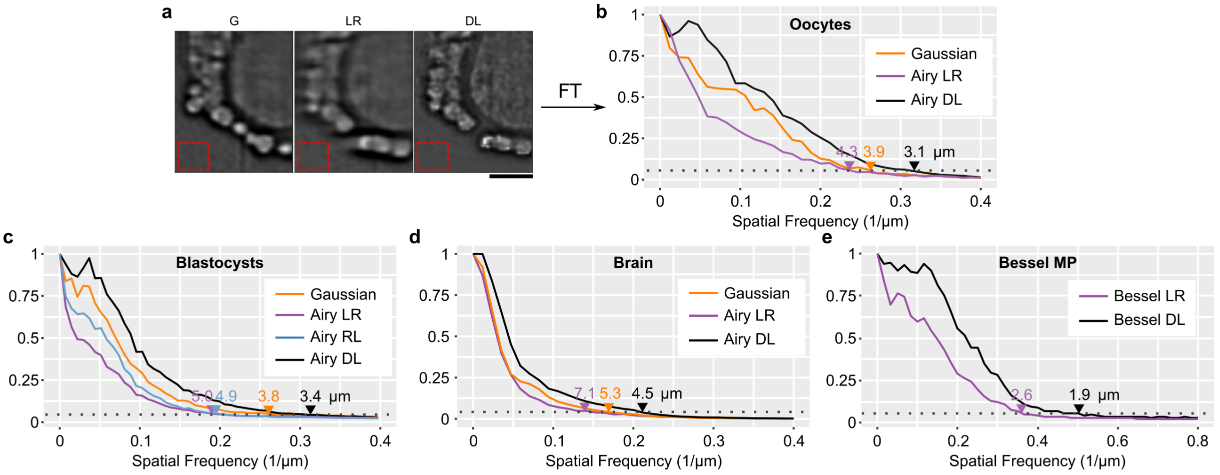


**Fig. S10.** Quantification of resolution in biological samples. (a) Filtered and cropped sample images. The red rectangle represents a region classified as noise for analysis. (b–e) Fourier transforms of filtered sample images approximating the modulation transfer function. The dotted lines represent the 5% cross-over value, approximating the FWHM resolution. Scale bar is 40 $\mu$m.

#### Supplementary Note 11: Widefield imaging of embryos

Figure S11 illustrates the widefield imaging capacity of Airy LSM. Figures S11 (a) and (b) visualise a 1-mm transverse FOV. Cross-sections marked by red dashed lines are visualised in Figu. S11(c) and (d), and support that data in Fig. 6. The insets in Figs S11 (a) and (b) illustrate an oocyte arrested at the metaphase II stage of meiosis. During metaphase, the chromosomes are aligned and held in place by the meiotic spindle along a plane termed the metaphase plate. This can be seen by the white line that separates two darker spherical regions that depict the spindle barrel. This morphology is particularly prominent in the Airy DL images, and can be confirmed for all oocytes by inspecting the volume (see Data Availability). This is an essential event for an oocyte undergoing nuclear maturation, necessary for fertilisation. As described for the blastocyst-stage embryo, the small bright regions within the oocyte are indicative of metabolic activity of mitochondria through FAD autofluorescence. The multitude of morphological features enhanced by Airy DL over a wide FOV and depth range underlines its potential for label-free and low-phototoxic imaging for embryo health and IVF success. Revealing the source and implications of these markers would be of considerable interest for future bioimaging studies.

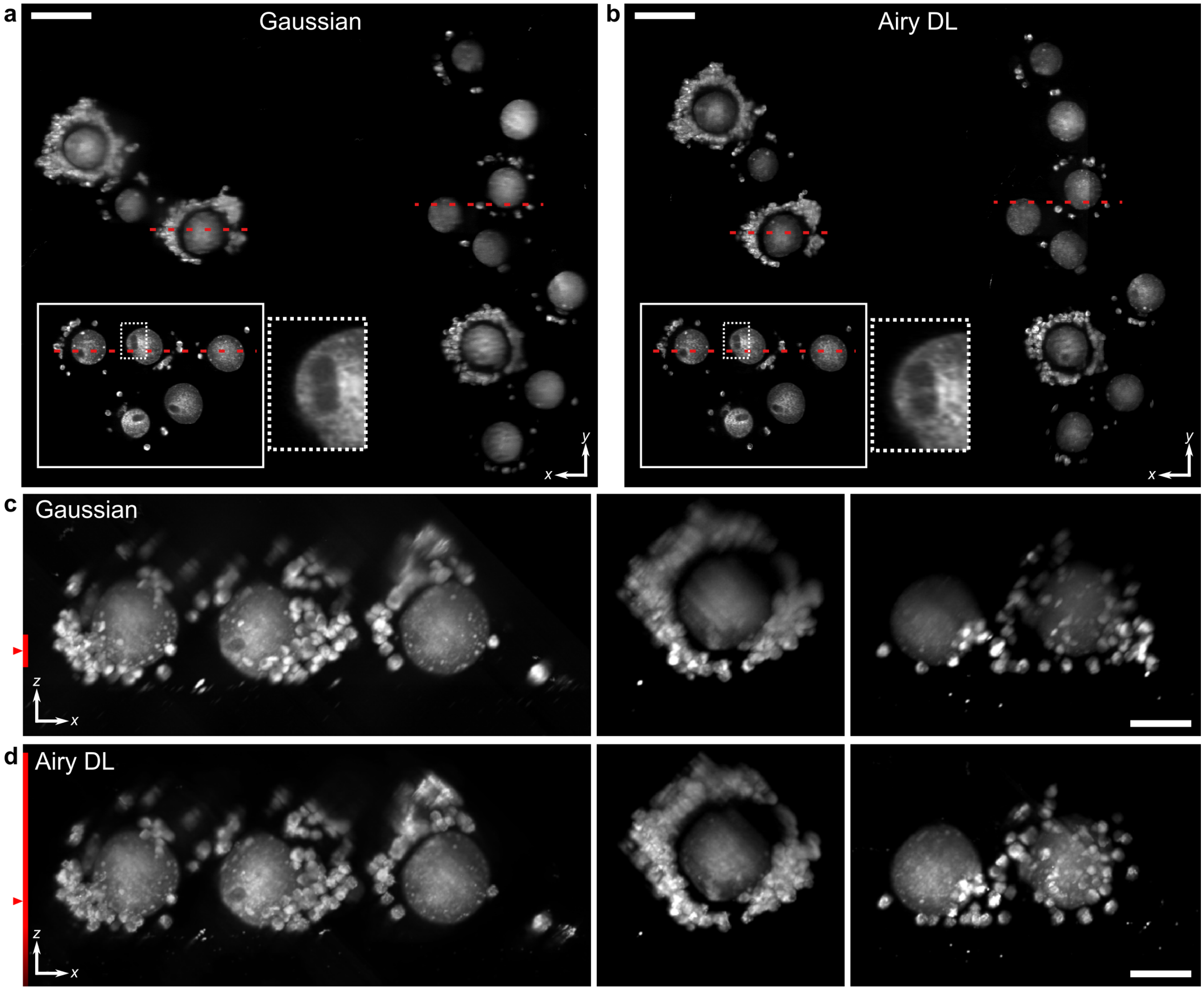


**Fig. S11.** Widefield 3D imaging of mouse cumulus oocyte complexes (COCs) with a Gaussian and Airy beams with deep learning deconvolution. (a,b) Enface sections of the Gaussian (a) and Airy (b) LSM images. The inset marked by a solid line in (a,b) is taken from another sample using identical imaging parameters. Scale bars in (a,b) are 100 $\mu$m. The zoomed-in region marked by a dashed line depicts the barrel-shaped spindle and metaphase plate of meiosis II. The red dashed lines mark the location of the cross-sections in (c,d). (c,d) Cross-sectional maximum intensity projections (range: 50 $\mu$m). The red line on the left of (c,d) shows the theoretical DOF of the Gaussian and Airy beams. The red triangle marks the focal position. Scale bars in (c,d) are 50 $\mu$m.

#### Supplementary Note 12: Application of learned deconvolution to open-source data: Bessel-beam LSM

Here, we demonstrate the versatility of our DL method by applying it to data generated by others. Specifically, on Medaka embryo timelapse images generated by Takanezawa *et al.,* (18). LR images were acquired using a multiphoton LSM with a Bessel beam (18). We deconvolved the maximum intensity projection of these timelapse volumes using the same DL network trained for the in-house build Bessel LSM (Fig. 7), but with a pixel resolution of 2.88 $\mu$m (compared to 0.85 $\mu$m in all other results). This pixel sampling was matched to the original data (18) and visually corresponded to the PSF. The pixel sampling of the original data was too coarse to capture the oscillating side-lobe structure of the Bessel beam, but the broadened intensity has remained. Figure S12 illustrates the performance of DL on maximum intensity projections from selected time points. It is evident in the insets that the resolution is substantially improved with DL. In Fig. S12(c), the line plots show the enhancement in resolution through selected regions marked by the red triangles. Using Fourier analysis (Supplementary Note 10), the resolution was quantified to be 11 $\mu$m in the LR images and 6.4 $\mu$m in DL images. The full timelapse videos are available as Supplementary Movies S1 and S2. The ease of training and application of the model suggests that our method may be readily translated to other imaging applications that use propagation-invariant beams.

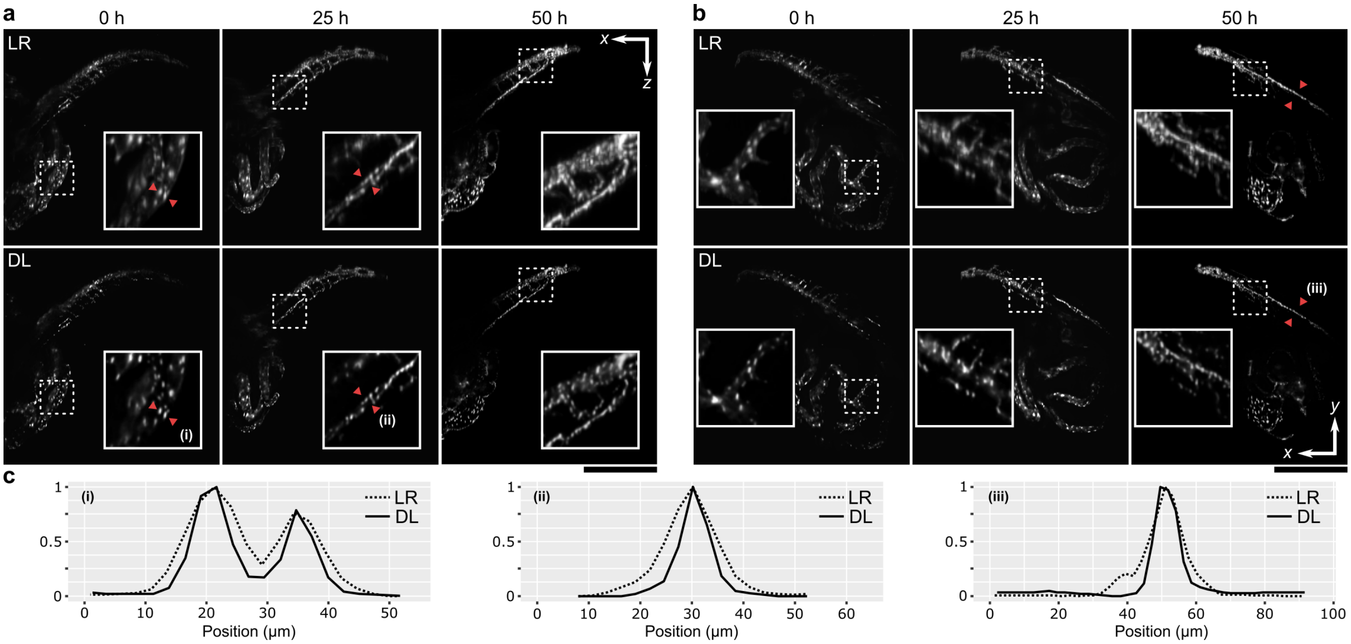


**Fig. S12.** Timelapse images of a developing Medaka embryo (78 hours past fertilisation at $t = 0$) with a multiphoton LSM with Bessel beam. Low resolution images were provided in (18) as open source. (a, b) Maximum intensity projections in the cross-sectional (a) and en-face (b) planes. Low resolution (LR) images are deconvolved (DL). Insets correspond to regions marked by the white rectangle.(c) Profile plots corresponding to locations marked by the red triangles. Scale bar is 500 $\mu$m.

#### Supplementary Note 13: Comparison of learned deconvolution with published networks

Here, we compare our work to the network and the training methodologies of a selection of contemporary state-of-the-art methods that have emerged in microscopy (19). Deep learning methods for image enhancement comprise two important features. First, is the network architecture, such as the arrangement and composition of network layers. Second, is the training methodology, such as how the training data is acquired or generated, and how it is used to provide strong losses that lead the network to effectively perform a desired transformation task. Arguably, it is the training methodology that has the most profound impact on deep learning methods and have led to the major advances in microscopy image enhancement. Early demonstrations, such as content-aware reconstruction (CARE) in microscopy (14) and cross-modality super-resolution (CMSR) (9), have shown that the choice and quality of training data led to effective image enhancement, tailored to tasks such as super-resolution (CARE Deconvolve), denoising in low-SNR imaging (CARE Denoise), or a reconstruction of an isotropic resolution, whilst using off-the-shelf U-Net architectures (20), initially proposed for image segmentation tasks. CMSR networks were trained using a U-Net with an added adversarial loss, initially proposed by Isola *et al.,* (5). The added adversarial loss in the GAN implementation led to improved training when low resolution and ground truth images featured different image content. Beyond U-Nets, the use of dense residual networks (ResNet) have demonstrated improved performance for image super-resolution and enhancement tasks (1, 2). In more recent work, Chen *et al.,* (15) have extended cross-modality image restoration with a three-dimensional residual channel attention network (RCAN), proposed by Zhang *et al.,* (21), for enhanced denoising and super-resolution, especially in low-SNR regimes. The RCAN uses a residual-in-residual architecture that enables more effective training when the network is very deep compared to ResNet-based networks (21). These methods are data-driven in that they rely on 1,000–10,000s carefully matched low-resolution images and their respective ground truths. As an alternative, Zhang *et al.,* (16) have proposed the use of artificially degraded low resolution images from experimental ground truths (DSP-Net Upscale), as well as computationally reconstructed ground truths using the SRRF algorithm (DSP-Net Content), to reduce the burden of image matching. This work employed a dual-stage processing network (DSP-Net), which featured a down-up-sampling layer set for image noise reduction before a more conventional ResNet architecture. Many methods have been proposed recently to reduce this data burden (13, 22). However, most methods have relied on either access to a high-resolution ground truth (to be degraded to a low resolution input pair) or another algorithm to computationally generate a ground truth.

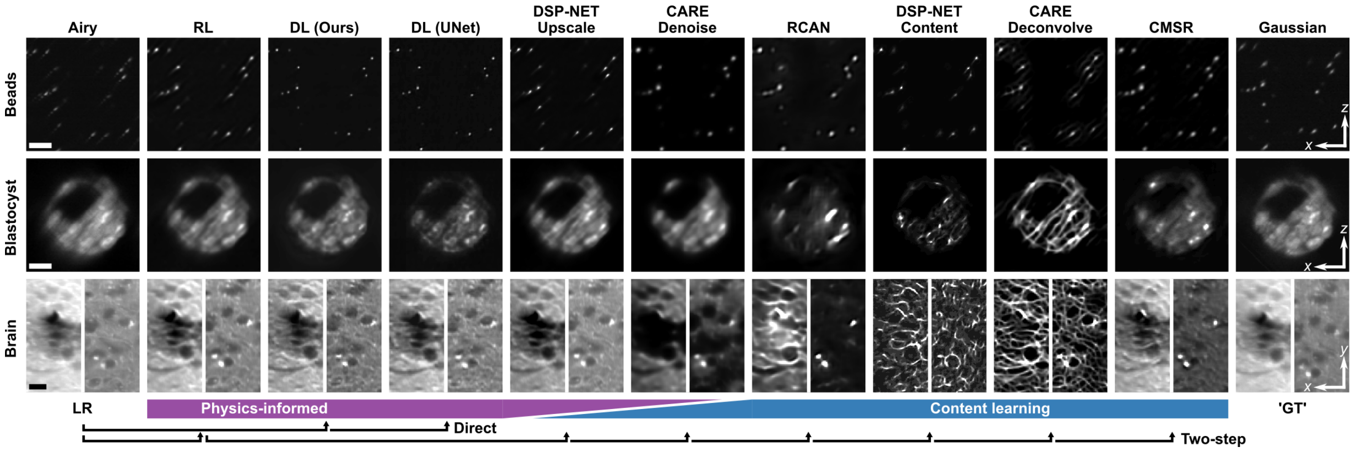


**Fig. S13.** Illustration of physics-informed learning in the context of contemporary deep learning microscopy methods. Scale bar: 20 $\mu$m.

To reconcile these impressive methods with the role of deconvolution in bioimaging we consider the following scenario. Specifically, the case where a user acquires a microscopy image or volume with a known PSF and desires the data to be deconvolved. This is the likely scenario for most application of bioimaging, and in this case, access to any ground truths or extensive experiments with super-resolution modalities is not available. Here, the options for deconvolution are to use conventional deconvolution, such as RL deconvolution or to incorporate a trained or 'demo' deep learning model.

In contrast to these methods, we propose that an image enhancement network can be trained without any measurement or estimate of a ground truth. Figure S13 provides a visual comparison of methods available in the instance where only a single volume of a low-resolution image is available and is sought to be deconvolved. Specifically, we demonstrate outputs of DSP-Net (16), CARE (14), RCAN (15) and CMSR (9) 'demo' models pre-trained on biological ground truths. To accommodate an Airy PSFs, we perform two-step reconstruction by performing Richardson-Lucy (RL) deconvolution on the Airy LSM image (LR) first and then processing the result using the relevant pre-trained or 'demo' network. Outputs are then compared to the same area images with a Gaussian beam as the ground truth (GT), and quantified using simulation in Table S1. It is possible to re-train the above networks with the same physics-consistent image pairs and loss functions proposed in this work. However, since many of the networks share very similar architectures, because performance is strongly dependent on training data, the output quality would likely be similar. Here, instead of comparing the utility of individual building blocks of neural networks, we seek to illustrate the main differences between physics-informed and content learning, which we will discuss presently. Our implementation of DL is similar to DSP-Net and RCAN in the use of a residual architecture. We also present outputs of our DL method trained using an alternative U-Net architecture (5) (DL (U-Net)) that underlies CMSR, CARE denoise and CARE deconvolve (Supplementary Note 4).

We can consider deep learning to comprise physical and content learning (8, 13). Physical learning encourages consistency with the transformation by the imaging system, for instance, the blur from the finite bandwidth of optical elements or the noise in detectors. Content learning encourages output images to resemble features observed in the ground truths related to, for instance, the type of samples or spectral contrast expected. The greatest constraint through the combination of physical and content learning, for example by training on a single sample type and instrument, has the potential to lead to the greatest performance gain whilst compromising generality. For instance, networks trained to super-resolve microtubules may demonstrate a high resolution and contrast enhancement, but will likely struggle to reconstruct images of other samples (19). Despite the performance of content learning, it is often desirable to achieve a generalised network that learns the physical inversion of the imaging system agnostic of the sample, such that one may be confident that the output is faithful to the imaged data. Physical learning may be encouraged by incorporating imaging physics in the network architecture or the loss functions, or by providing training images that emphasise physics transformations (22).

Table S1 quantifies the performance of each method using simulated data based on beads and biological images based on peak signal-to-noise ratio (PSNR) and resolution (Supplementary Note 10). It is immediately evident that those networks that have learned indiscriminately the physics and content of paired images (marked as content learning in Fig. S13) have vastly varied performance across sample type that is often inconsistent with the expected GT. Specifically, RCAN was trained on labelled endoplasmic reticulum in U2OS cells, DSP-Net Content on labelled microtubules in U2OS cells, and CARE Deconvolve on microtubules in INS-1 cells. In these networks, outputs resemble the style of the training set regardless of the input image. DSP-Net Content outputs present the highest resolution, but are visually inconsistent with the GT. The CMSR network was trained on TxRed labelled endothelial cells and presents improved generality across sample types. However, the relative intensity in the images emphasise bright features over the GT. We note that this is common to U-Net architectures (Supplementary Note 4), which can lead to greater potential for improved resolution at the expense of a reduction in PSNR in dense tissues, as evident by comparing the performance of our DL method with DL (U-Net). The CARE Denoise network was trained on high- and low-SNR acquisitions of the sample sample and system. This training favoured system noise reduction instead of image content transformation, which is evident by the low resolution of its outputs. However, when provided with good SNR images, the network has attenuated regions of low image intensity, which is especially evident in the brain. DSP-Net Upscale was trained on high-resolution GTs that have been digitally downsampled with added noise. Thus, the training favoured the physical transformation of upscaling and denoising. While DSP-Net Upscale demonstrates outputs consistent with the GT, the outputs offer little quantitative improvement over RL deconvolution. In contrast, our method (DL), which is designed for deconvolution, offers superior visual contrast and resolution in all sample types. When we retrained our method using a U-Net (DL U-Net), while presenting improved resolution, the PSNR of dense biological tissues was substantially lower. This is likely due to spatial sparsity being favoured by U-Nets because of its autoencoder design (Supplementary Note 4). We employ the ResNet DL architecture for the demonstrations in this work because of its high PSNR and low computational demand. Independent from the network architecture, however, the major distinction of our method to those published previously is that our learning approach is free from any experimental ground truth.

| | **RL** | **DL (Ours)** | **DL (UNet)** | **DSP Upscale** | **CARE Denoise** | **RCAN** | **DSP Content** | **CARE Deconv.** | **CMSR** | **GT** |
|---|---|---|---|---|---|---|---|---|---|---|
| **PSNR Bead (dB)** | 33.0 | **38.6** | 37.3 | 21.2 | 27.5 | 25.4 | 22.4 | 29.1 | 30.8 | |
| **Res. Bead (um)** | 3.1 | 2.0 | **1.9** | 2.8 | 4.1 | 5.7 | 2.1 | 3.2 | 2.4 | 2.6 |
| **PSNR Bio (dB)** | 30.9 | **34.0** | 28.7 | 30.3 | 27.2 | 21.2 | 20.4 | 23.3 | 25.0 | |
| **Res. Bio (um)** | 3.7 | 2.8 | 2.6 | 2.8 | 14.6 | 4.4 | **2.0** | 3.2 | 2.6 | 4.0 |
| **Architecture** | | ResNet | UNet | ResNet* | UNet | RCAN | ResNet* | UNet | UNet | |
| **GPU RAM (GB)** | | 6 | 6 | 11 | 8 | 12 | 11 | 8 | 8 | |
| **Data Size (MPs)** | | 4† | 4† | 580 | 1000 | 1000 | 200 | 90 | 2700 | |
| **Training Time** | | 1.5h | 1.5h | 14h | 30h | 50h | 5h | 1.5h | 10–20h | |
| **Inference** | 0.2–1s | 0.2s | 0.2s | 1s | 0.2s | 4s | 1s | 0.2s | 0.4s | |

**Table S1.** Comparison of deep learning methods. Deconvolution is quantified using the peak signal-to-noise ratio (PSNR) and resolution (Res.) metrics. GPU RAM is based on the maximum RAM available during training for the respective methods. Training and inference time are estimated based on 1 megapixel image sizes and a state-of-the-art GPU. Notes: (*) DSP-Net utilised ResNet blocks with an extra denoising layer; (†) Data pairs are simulated and not experimental.

For a further comparison we have listed additional performance and computational load metrics. The GPU RAM usage by the networks is estimated based on the maximum RAM of the GPU used in the respective publications, which has implications on the maximum network size and the reported speed of training. The data size is presented in a normalised fashion as the total pixel count of the GT images. Because of the relatively small network and simulated data, our DL approach has a substantially lower data burden compared to other methods. This is also because our network does not prioritise the content learning, which often requires large datasets, in contrast to cross-modality methods, such as RCAN and CMSR. The training and inference time is also low and similar to CARE deconvolution, and approach the lower bound of the time needed to transfer image data from disk to the GPU and back.

It is likely that applying our physics-informed training data and loss functions to more sophisticated networks, such as DSP-Net and RCAN, which include extra denoising steps, 3D image handling and deeper residual layers, would lead to improved overall performance, however, at the expense of training and inference time. This would be of interest for future studies, especially if the networks are to be trained once per system based on the high generality.

#### Supplementary Note 14: Structured light fields

In this section, we describe the formation of the Gaussian, Airy and Bessel beams through the control of the light field intensity and phase in the pupil plane. For consistency, we define the electric fields at the pupil and image planes as $E_p$ and $E_i$, respectively. Further, for simplicity, we describe each beam in 2D, with respect to the transverse ($x$) and depth ($z$) coordinates. This can be trivially extended to 3D through the rotational symmetry of the Gaussian and Bessel beams, and through a multiplication of the Airy profiles in $x$ and $y$. We note that with the SLM, we are able to control the intensity not amplitude, thus, we square the desired amplitude prior to projection.

**Gaussian beam.** A Gaussian beam in the image plane can be described by the Gaussian function:

$$E_i(x) = \exp\left(\frac{-x^2}{w_i^2}\right), \qquad \textbf{(1)}$$

where $w_i$ is the $1/e$ radius of the amplitude ($1/e^2$ radius of intensity). Using the Fourier-transforming property of a lens, we can define the field at the pupil plane (with a normalised amplitude) as:

$$E_p(x) = \exp\left(\frac{-x^2}{w_p^2}\right), \qquad \textbf{(2)}$$

where $w_p = f\lambda/(\pi w_i)$. Thus, we can trivially generate a Gaussian beam at focus by projecting a Gaussian intensity onto the SLM.

Gaussian beam propagation is well-known, and given as:

$$E_i(x,z) = \frac{w_i}{w(z)} \exp\left(\frac{-x^2}{w(z)^2}\right) \exp\left(-i(kz + k\frac{x^2}{2R(z)} - \psi(z))\right), \qquad \textbf{(3)}$$

where $w(z) = w_i\sqrt{1+(z/z_R)^2}$, $z_R = \pi w_i^2 n/\lambda$ is the Reyleigh range, $R(z) = z(1+(z_R/z)^2)$ is the radius of curvature, $\psi(z) = \arctan(z/z_R)$ is the Guoy phase, and $n$ is the refractive index.

**Airy beam.** The Airy beam is a solution to the paraxial equation of diffraction in 1D, given as (23, 24):

$$E_i(\xi, s) = \mathrm{Ai}(s - (\xi/2)^2)\exp(i(s\xi/2) - i(\xi^3/12)), \qquad \textbf{(4)}$$

where Ai is the Airy function, $s = x/x_0$, $\xi = z/kx_0^2$, $k = 2\pi n/\lambda_0$, $x_0$ is a dimensionless coordinate. Such an Airy beam is 'non-diffracting', however, it has infinite energy. A finite-energy Airy field can be realised by apodising the intensity in the cross-section by an exponent $\exp(\gamma s)$, resulting in the following form:

$$E_i(\xi, s) = \mathrm{Ai}(s - (\xi/2)^2 + i\gamma\xi)\exp(\gamma s - \gamma\xi^2/2 - i\xi^3/12 + i\gamma^2\xi/2 + is\xi/2). \qquad \textbf{(5)}$$

We make an important note here that in (23, 24) the apodisation parameter was termed as $\alpha$. We instead term this parameter as $\gamma$ to avoid ambiguity with the Airy $\alpha$ parameter that scales the cubic phase that we will encounter in the pupil plane. This is to be consistent with previous publications (25–27).

In the image plane (at focus, $\xi = 0$), the finite-energy Airy field is given as:

$$E_i(x) = \mathrm{Ai}(s)\exp(\gamma s). \qquad \textbf{(6)}$$

The field at the pupil plane is related to the Fourier transform:

$$E_p(s_2) = \exp(-\gamma s_2^2)\exp(is_2^3/3 - i\gamma^2 s_2 + \gamma^3/3), \qquad \textbf{(7)}$$

where $s_2$ is evaluated at lens coordinates $s_2 = s/(f\lambda)$. We recognise that the field comprises a Gaussian intensity and a phase profile that is dominated by a cubic term. We can approximate this as:

$$E_p(s_2) = \exp(-\gamma s_2^2)\exp(is_2^3/3). \qquad \textbf{(8)}$$

We can substitute more intuitive parameters, such that:

$$E_p(x) = \exp(-x^2/\omega_p^2)\exp(i(2\pi\alpha/\omega_p^3)x^3) \qquad \textbf{(9)}$$

where $\alpha$ indicates the number of phase wraps at $\omega_p$. These pupil parameters relate to the Airy solution in Eq. (5) via: $\gamma = 1/(6\pi\alpha)^{2/3}$ and $x_0 = \lambda f(6\pi\gamma)^{1/3}/(2\pi\omega_p)$, which enable the modelling of Airy propagation at focus. We note that in previous publications (25–27), $\alpha$ parameters were arbitrarily scaled in the pupil coordinate space. Our representation is scale-invariant, such that $\gamma$ is solely dependent on $\alpha$, and $x_0$ is solely dependent on $w_p$.

**Bessel beam.** The Bessel beam, described by the Bessel function along the radial coordinate, is a non-diffracting beam solution to the wave equation. Here, we only consider a zero-order Bessel beam, which is described by:

$$E(x,z) = J_0(k_x x)\exp(ik_z z)\,, \tag{10}$$

where $k_x$ and $k_z$ are transverse and longitudinal wave vectors: $k = \sqrt{k_x^2 + k_z^2} = 2\pi/\lambda$, and $J_0$ is the zero-order Bessel function. A finite-energy field can be described by a Bessel modulated by a Gaussian envelope—a Bessel-Gauss beam, described in the focus ($z = 0$) as:

$$E_i(x) = J_0(k_x x)\exp(-x^2/w_i^2)\,, \tag{11}$$

where $w_i$ is the waist of the Gaussian modulation beam.

The pupil function can be described by a Fourier transform of the beam at focus. Following the derivation in (28), using the Bessel function identity, the pupil function (with normalised amplitude) is:

$$E_p(x) = \exp\left(-\frac{x^2 + r_r^2}{w_p^2}\right) I_0\left(\frac{2r_r x}{w_p^2}\right)\,, \tag{12}$$

where $w_p = 2f/(kw_i)$, $r_r = k_x f/k$ and $I_0$ is the modified Bessel function of zero order and of first kind.

Typically, $r_r$ is much larger than $w_p$. In this case, $I_0(x) \sim \exp(x)$, reducing the above equation to:

$$E_p(x) = \exp\left(-\frac{(x - r_r)^2}{w_p^2}\right)\,. \tag{13}$$

This describes a ring with a Gaussian thickness in the pupil plane.

The propagation of the Bessel-Gauss beam at focus can be exaluated based on the work of (29). As the field propagates in $z$, it is subject to diffraction, which can be described by the Fresnel diffraction integral:

$$\begin{aligned} E_i(x,z) = {} & (-ik/z)\exp(i(kz + kx^2/(2z))) \\ & \times \int_0^\infty E(x_0,0)\exp(ikx_0^2/(2z))J_0(kx_0x/z)x_0 dx_0\,, \end{aligned}$$

which can be transformed into the following form (29):

$$\begin{aligned} E_i(x,z) = {} & (w_i/w(z)) \\ & \times \exp\{i[(k - k_x^2/(2k))z - \Phi(z)]\}\, J_0(k_x x/(1 + iz/L)) \\ & \times \exp\{[-1/w(z)^2 + ik/(2R(z))](x^2 + k_x^2 z^2/k^2)\}\,, \end{aligned}$$

where

$$\begin{aligned} L &= kw_i^2/2 \\ w(z) &= w_i[1 + (z/L)^2]^{1/2} \\ \Phi(z) &= \arctan(z/L) \\ R(z) &= z + L^2/z \end{aligned}$$

We note the similarities to Gaussian beam propagation, where $R(z)$ is the radius of curvature, $\Phi(z)$ is the Gouy phase, $w(z)$ is the spot size parameter, and $L$ is an effective analog of the Rayleigh range. All parameters can be evaluated from pupil paramters: $w_i = 2f/(kw_p)$ and $k_x = r_r f/k$.

#### Supplementary Note 15: Supplementary Movies

**Supplementary Movie S1** is a maximum intensity projection in the $xz$ plane of the timelapse in Fig. S11.
**Supplementary Movie S2** is a maximum intensity projection in the $xy$ plane of the timelapse in Fig. S11.
